# Recurrent evolution of two competing haplotypes in an insect DNA virus

**DOI:** 10.1101/2020.05.14.096024

**Authors:** Tom Hill, Robert L. Unckless

**Affiliations:** 4055 Haworth Hall, The Department of Molecular Biosciences, University of Kansas, 1200 Sunnyside Avenue, Lawrence, KS 66045

**Keywords:** nudivirus, *Drosophila innubila*, DiNV, virulence, coevolution

## Abstract

Hosts and viruses are constantly evolving in response to each other: as hosts attempt to suppress the virus, the virus attempts to evade and suppress the host’s immune system. This arms race results in the evolution of novel pathways in both the host and virus to gain the upper hand. Here we describe the coevolution between *Drosophila* species and a common and virulent DNA virus. We identify two distinct viral types that differ 100-fold in viral titer in infected individuals, with similar effects across multiple species. Our analysis suggests that one of the viral types appears to have recurrently evolved at least 4 times in the past ∼30,000 years, including in another geographically distinct species, due to the high effective mutation rate which increases with titer. The higher titer viral type is associated with suppression of the host immune system and an increased transmission rate compared to the low viral titer type. Both types are maintained in all populations, likely due to an increased virulence in the high titer type creating a trade-off between effective transmission and virulence and resulting in nearly equal reproduction rates (R_0_) in both types. Together these results suggest that the reciprocal selective pressures caused by the co-evolution between host and virus has resulted in this recurrently evolving relationship.

## Introduction

Antagonistic coevolution between hosts and their parasites is nearly ubiquitous across the tree of life. As a result, genes involved in immune defense are among the fastest evolving genes in host genomes (Nielsen *et al*. 2005; Sackton *et al*. 2007; Enard *et al*. 2016; Shultz and Sackton 2019). Viruses are a particular fitness burden on hosts; for viruses to persist within populations, they must successfully invade the host organism, contend with the host immune system, replicate and then transmit the newly produced particles to a new host (Holmes 2007; Gifford 2012). Once successfully established in a population, natural selection acts to modulate the rate the virus propagates relative to its virulence, optimizing the ratio of viral virulence to transmission (Williams and Nesse 1991; May and Nowak 1995; Lipsitch *et al*. 1996). Thanks to their elevated mutation rate and high population sizes, viruses can easily do all of this, and even co-opt or manipulate host-pathways in the process (Burgyan and Havelda 2011; Davey *et al*. 2011; Palmer *et al*. 2019). As a result, proteins that interact with viruses often show the fastest rates of evolution, and the highest rates of adaptation compared to the rest of the genome, as the host attempts to suppress the pathogen, or escape infection (Obbard *et al*. 2006; Obbard *et al*. 2009a; Mukherjee *et al*. 2013; Enard *et al*. 2016; Palmer *et al*. 2018a).

Hosts have evolved numerous immune pathways to reduce viral burden and enhance survival after infection (Merkling and Van RIJ 2013; West and Silverman 2018). Interestingly, several of these pathways (IMD, Toll, JAK-STAT) are generally associated with other pathogens, yet also respond to infection by viruses, though the specific mechanisms are not yet known (Hoffmann 2003; Hultmark 2003; Coccia *et al*. 2004; Zambon *et al*. 2005; Costa *et al*. 2009; Ferreira *et al*. 2014; West and Silverman 2018). In *Drosophila melanogaster*, the RNA interference (RNAi) pathway is involved in resisting viral infection by generating suppressive RNAs complementary to the viral sequence (Hultmark 2003). As might be expected, this antiviral RNAi pathway is also rapidly evolving in many species (Obbard *et al*. 2006; Wang *et al*. 2006; Obbard *et al*. 2009b; Merkling and Van RIJ 2013).

Drosophila innubila Nudivirus (DiNV), which infects the host *D. innubila*, is the one of the few DNA viruses to naturally infect *Drosophila*, and the only documented case of a DNA virus infection at high frequency in natural *Drosophila* populations (Unckless 2011; Hill *et al*. 2019). *Drosophila innubila* is a mushroom feeding species of the *Drosophila* subgenus found inhabiting woodlands found on mountains across Arizona & New Mexico, separated by large expanses of desert. *D. innubila* radiated north from Mexico during the last glaciation period and came to inhabit these forests after the glacial retreat, creating a subdivided population with high rates of gene flow between locations (Dyer and Jaenike 2005; Hill and Unckless 2020).

During this period *D. innubila* likely became infected with DiNV, suggesting a long-lasting host-pathogen relationship in multiple populations. This could lead to opportunities to study the coevolution of DiNV and *D. innubila* in replicate (Hill and Unckless 2020), which could potentially result in parallel or divergent evolution of the virus and interacting host pathways (Anderson and May 1982; Kaltz and Shykoff 1998).

A pair of previous studies examined the rates of evolution of DiNV and *D. innubila*, finding the envelope and replication machinery to be rapidly evolving in DiNV, suggesting its importance in viral propagation (Hill and Unckless 2018). In *D. innubila*, the antiviral RNAi machinery is not rapidly evolving, possibly as DNA viruses interact with different immune pathways to RNA viruses (the primary burden of *D. melanogaster*) (Webster *et al*. 2015). Consistent with this, the Toll pathway is both rapidly evolving in *D. innubila* and suppressed by a related nudivirus upon infection in *D. melanogaster* (Hill *et al*. 2019; Palmer *et al*. 2019).

DNA viruses, and nudiviruses such as DiNV in particular, have much larger genomes (100kbp or larger) than RNA viruses, with much more complicated replication cycles (Rohrmann 2013). RNA viruses also have much higher mutation rates than DNA viruses, yet much lower levels of diversity due to lower recombination rates and efficient selection on variation in the genome (Pennings 2012; Pennings *et al*. 2014; Wilson *et al*. 2016; Feder *et al*. 2019). As a result, we expect the evolutionary dynamics of DNA viruses to differ dramatically from RNA viruses (Rohrmann 2013; Hill and Unckless 2017). In fact, because of their large genomes, high recombination rate and low mutation rate, we expect that DNA virus coevolution with hosts will be qualitatively different than RNA viruses or bacteria. Together this paints a picture suggesting that hosts can have different relationships with different pathogens (such as RNA viruses or DNA viruses), and the pathogens themselves can behave differently within the host. Characterizing the relationships between different species and their long-term pathogens, such as DiNV and *D. innubila*, will help broaden and expand our understanding of how hosts and pathogens evolve in response to each other.

Here, we survey the genetic variation in DiNV to infer its co-evolutionary history with *D. innubila* and two other associated hosts. We identify two viral multilocus genotypes (considered to be haplotypes) that differ by 11 focal SNPs and show that these viral types are maintained within the same host population and across multiple isolated host populations. Despite high rates of recombination, these SNPs are tightly linked likely due to extremely strong selection and possibly incompatibilities between types. One viral type is associated with 100-fold higher viral titer and increased virulence compared to the other. Further, we find evidence that the high titer type evolved independently in at least four geographically-isolated host populations. Together, these results suggest rapid evolutionary dynamics of host-virus interactions, due to the multiple competing viral types that interact with different host pathways.

## Results

### DiNV segregates for linked variants strongly associated with viral titer

To characterize the evolutionary dynamics of wild Drosophila innubila Nudivirus (DiNV) in its host (*D. innubila*), we sequenced wild-caught individuals from four populations with the expectation that some (∼40% in previous samples) individuals would be infected (Hill and Unckless 2020). We considered strains to be infected with DiNV if they had at least 10x coverage for 95% of the genome. In total, we used sequencing information for 57, 92, 92, and 92 individuals from the Huachucas (HU), Santa Ritas (SR), Chiricahuas (CH), and Prescott (PR) populations with infection rates 26%, 44%, 63% and 79%, respectively (Supplementary Table 1). We also used 35 individuals collected in the Chiricahuas in 2001 (52% infected with DiNV) and 80 individuals collected in the Chiricahua’s in 2018 (Supplementary Table 2, pre-selected using PCR, 40 infected with DiNV and 40 uninfected).

We isolated and sequenced DNA from these samples and, after filtering and mapping to the genome we called variation in the viral genomes to assess the extent of adaptation in each viral population. Consistent with an arms race between host and virus, most envelope and novel virulence (GrBNV-like) genes show strong signatures of adaptive evolution in each population compared to background viral genes (using McDonald-Kreitman based statistic Direction of Selection (Stoletzki and EYRE-WALKER 2011) and Selection Effect (Eilertson *et al*. 2012), Supplementary Figure 1, DoS > 0, GLM *p*-value < 0.05).

Given these potential signatures of an arms race between *D. innubila* and DINV, we first attempted to determine if any host or viral genetic variation is associated with within-host viral titer, which we use here as a measure of virulence. For each virus-infected individual, we quantified viral titer (as viral genome coverage normalized to host autosomal genome coverage) and identified both host and viral polymorphisms. We then performed an association study across both host and virus variable sites to identify variants significantly associated with viral titer using PLINK (Purcell *et al*. 2007).

Of 5,283 viral SNPs in the 155kbp DiNV genome, 1,403 SNPs are segregating in at least 5 infected host individuals. Of those 1,403 SNPs, 78 are significantly associated with viral titer after multiple testing correction (FDR < 0.01, Figure 1A). Of these, 16 are within 2000bp of the start site of a gene, 18 are coding nonsynonymous, 11 are coding synonymous and 33 are intergenic. The most significantly associated SNP is a non-synonymous polymorphism in the active site of *Helicase-2* (Figure 1A, Supplementary Figure 2). The *Helicase-2* polymorphism is the only significantly associated polymorphism found segregating at a range of frequencies *within* individuals (Supplementary Figure 2). The frequency of this derived polymorphism has a negative relationship between viral titer and the derived SNPs frequency (GLM t-value = −20.516, *p*-value = 5.55e-62). However, when ranking samples by viral titer, the SNP frequency does not fit this expectation and several samples fixed for the ancestral allele also have a lower viral titer (Supplementary Figure 2).

**Figure 1:**
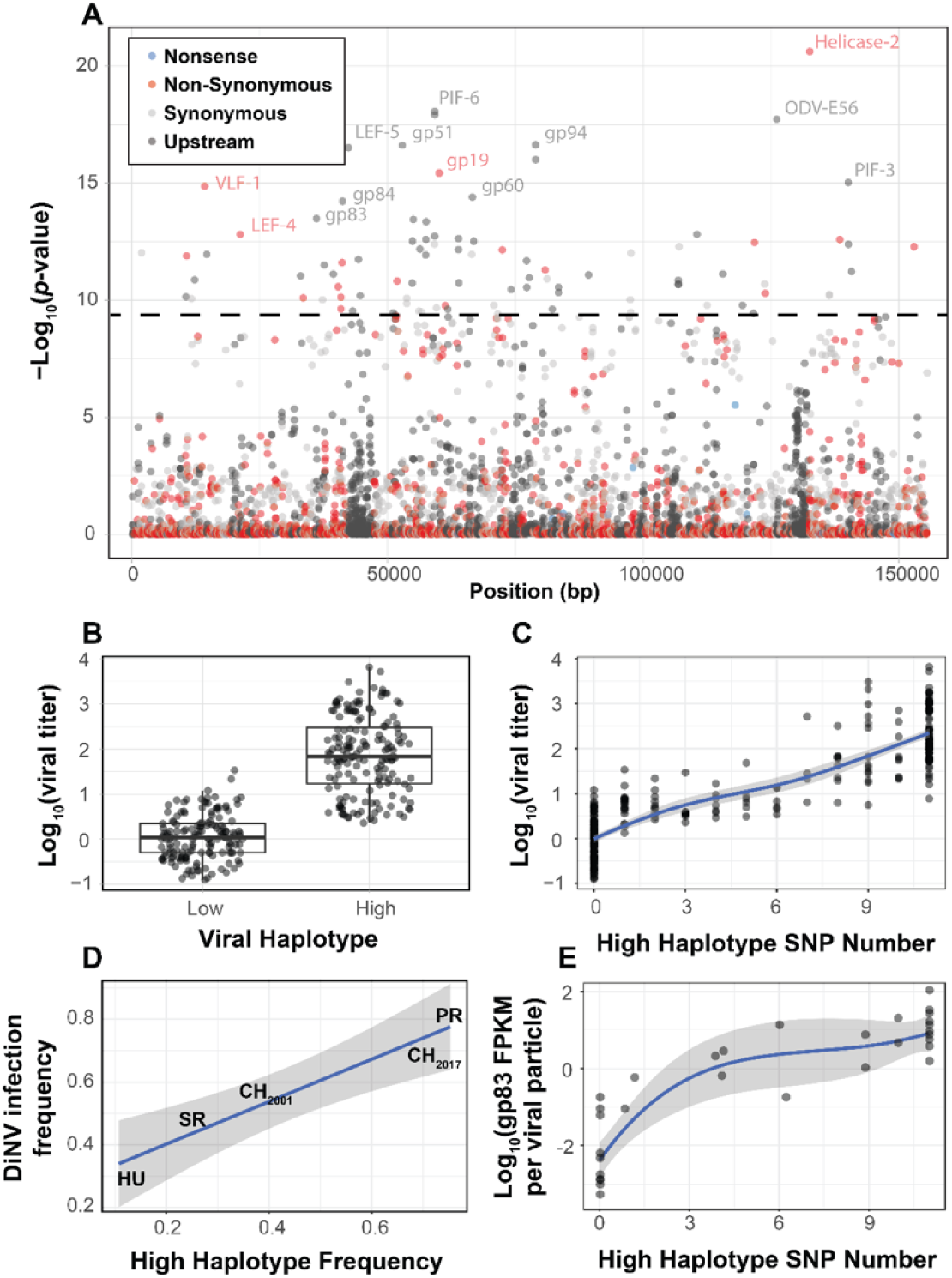
Viral genome-wide association study for DiNV titer in wild *D. innubila*. **A**. Manhattan plot for each DiNV SNP and the significance of its association with DiNV titer. SNPs are colored if they are upstream, synonymous, non-synonymous or nonsense mutations. 12 named SNPs are either part of the significantly associated viral haplotype or are in *Helicase-2*. The FDR corrected *p*-value cutoff of 0.01 is shown as a dashed line (multiple testing correction for 1,403 tests). **B**. Viral titer for individual wild caught flies infected with Low and High DiNV haplotypes (containing all high-type SNPs). The middle bar represents median value, upper and lower bars represent 25^th^ and 75^th^ percentile and whiskers represent a 95% confidence interval. **C**. Association between the number of significant SNPs and the viral titer of a sample. **D**. Across five populations, the frequency of the High DiNV haplotype is correlated with the frequency of the virus infection. **E**. Expression (in FPKM per viral particle) of *gp83* increases with the number of High DiNV haplotype SNPs.

We also identified a striking association between viral titer and eleven nearly perfectly linked polymorphisms found across the DiNV genome (Figure 1A, highlighted SNPs, Table 1, Supplementary Figures 2-4, Sig. SNPs). We refer to these two types as the ‘High Type’ and ‘Low Type’ (Figure 1B). This multilocus genotype includes three non-synonymous SNPs and five SNPs in the UTRs of known virulence factor genes and three intergenic SNPs. Viral titer is, on average, 100-fold higher in individuals infected with High type virus compared to the ancestral Low type (Figure 1B). Though we found few strains with an intermediate number of SNPs, viral titer increases as the number of High type SNPs increases (Figure 1C, GLM t-value = 34.971, *p*-value = 5.912e-16), though the rate of increase slows as the number of High type SNPs increases suggesting diminishing returns (Figure 1C). Some of these polymorphisms are associated with known virulence factors, or are related to the formation of the viral envelope co-opting the host vesicle trafficking system and are rapidly evolving in nudiviruses (e.g. *VLF-1, ODV-E56, PIF-3*) (Rohrmann 2013; Hill and Unckless 2017; Hill and Unckless 2018). Additionally, several are associated with orphan genes thought to be novel virulence factors, including a *gp83*, a gene that downregulates Toll-induced antimicrobial peptides (AMPs) and upregulates those induced by IMD (*gp83*) (Palmer *et al*. 2019). Both pathways may interact with DNA viruses (Zambon *et al*. 2005; Costa *et al*. 2009; Merkling and Van RIJ 2013; Ferreira *et al*. 2014; Lamiable *et al*. 2016; Palmer *et al*. 2019).

**Table 1:**
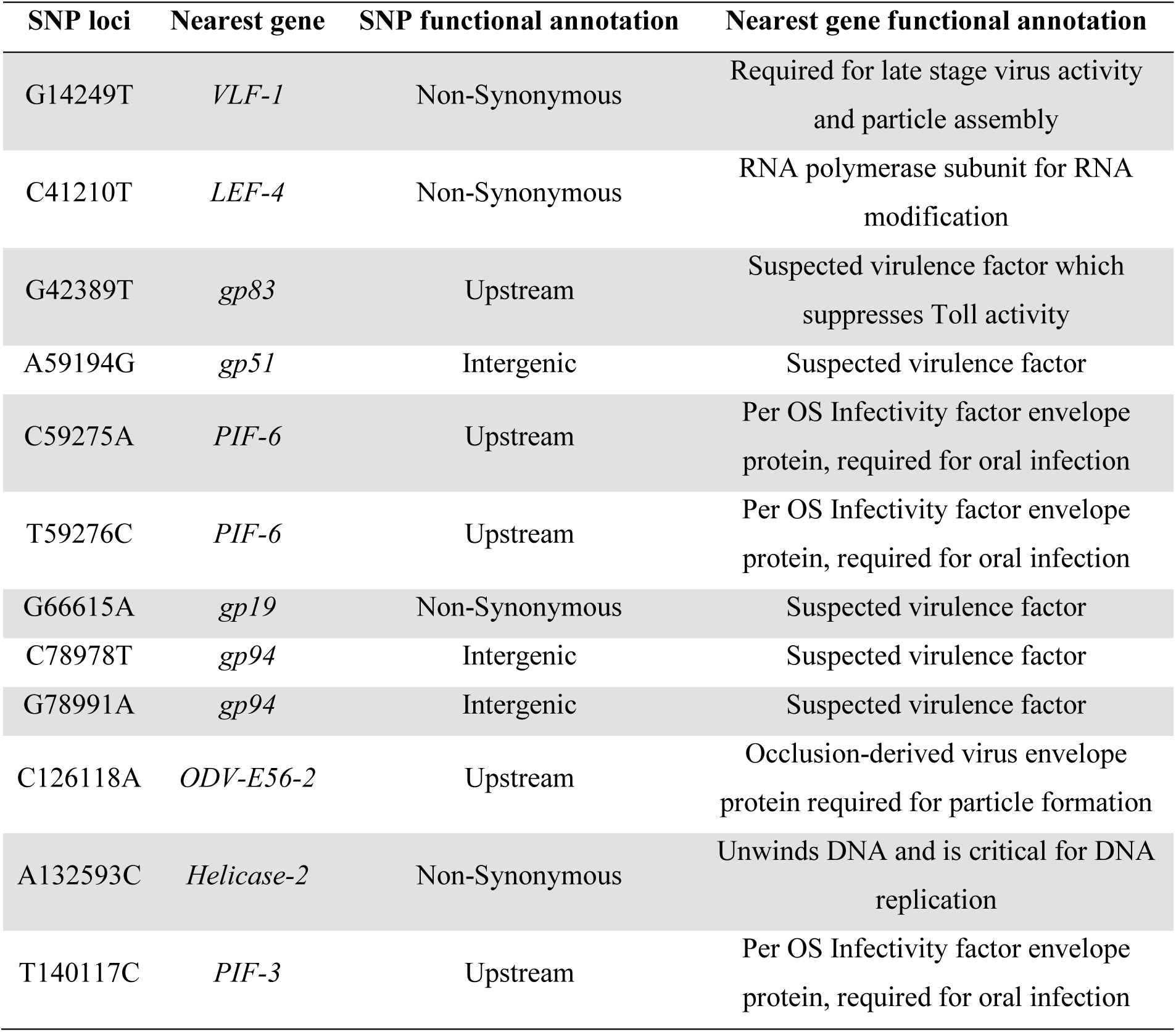
Candidate viral SNPs associated with viral titer, their loci, associated genes and the functional category of that gene.

Among populations there is a positive correlation between the frequency of the High type and overall DiNV infection frequency (Figure 1D, GLM logistic regression z-value = 6.104, *p*-value = 0.00883), suggesting that the High type may have a higher effective transmission rate, resulting in a higher number of new individuals infected, per DiNV infected individual. The transmission rate appears to be higher for the High type, despite a possibly higher death rate than the Low type. We also find both viral types in collections from 2001 and 2017, with the High type significantly more common in the 2017 collection (Fisher Exact Test *p*-value = 0.0167, Figure 1D).

Further, the expression of the virally encoded suppressor of Toll signaling, *gp83*, per viral particle is greater in strains containing at least one derived SNP of the high type increase (Figure 1E, GLM t-value = 10.32, *p*-value = 7.34e-12), suggesting enhanced virulence in the high type. For these SNPs, few are found at intermediate frequencies within samples (Supplementary Figure 2), and no samples contain more than two SNPs at a similar frequency (Supplementary Figure 2). This suggests that hosts are either infected completely with Low type or High type virus particles and the two types may be partially incompatible with each other. Together, these results suggest that the high viral type is more virulent possibly because it is better able to suppress host Toll signaling.

### The High DiNV viral type suppresses the Drosophila immune system and has elevated virulence

Given the striking difference in viral titer and infection frequencies across populations between High and Low type viruses, we sought to further characterize the differences in infection dynamics between the types. We sequenced mRNA from 80 wild *D. innubila* males collected in 2018 (Supplementary Table 2, 40 infected with DiNV, 40 uninfected) and performed a differential expression analysis between infected and uninfected individuals. Few genes were differentially expressed (DE) between infection states in *D. innubila* (Supplementary Figure 5A, *p-*value < 0.01 after FDR multiple testing correction, above the dotted line), but these DE genes were enriched for several interesting categories. Specifically, we found IMD-induced antimicrobial peptides (AMPs) were upregulated upon DiNV infection, while one Toll-induced AMP, and several chorion genes and heat shock proteins were downregulated (Supplementary Figure 5A). We also compared these results to a laboratory experiment in *D. melanogaster* of differential expression after infection with a close relative of DiNV (Palmer *et al*. 2018b). These same genes are also differentially expressed in *D. melanogaster*. Of the 12 genes which are differentially expressed in the same direction in both species, five are AMPs and five are chorion genes (Supplementary Figure 5B).

We then compared gene expression between *D. innubila* infected with High or Low type. We find 17 DE host genes and 9 DE viral genes between types (after controlling for virus copy number as FPKM/titer, Figure 2, FDR corrected *p*-value < 0.01, GLM t-value = −4.6239413, *p*-value = 9.876143e-05). Specifically, four Toll-mediated immune peptides (*Listericin, IM33*, Bomanins *BomBC2* and *BomT2*) have reduced expression in High Type infected individuals compared to the Low Type (Figure 2, Supplementary Figure 6). Finally, viral genes of interest (*PIF-3, VLF-1, gp83*) have higher expression per viral particle (FPKM/titer) in the High type compared to the Low type, all also increase in expression per viral particle (FPKM/titer) as the number of High type alleles increases (Figure 1E & 2, t-value = 13.732, *p*-value 3.36e-15). This pattern may be driven by the allele of the non-synonymous SNP in *VLF-1* (GLM t-value = 2.13, *p*-value = 0.04272) and the alleles of the SNPs upstream of *gp83, PIF-3, gp51* and *ODV-E56* (GLM t-value = 3.518, *p*-value = 0.00162). Together these results suggest that the high viral type has increased expression of key virulence factors, which in turn, manipulate the expression of host genes involved in immune defense to result in the observed differences in viral titer. These results suggest that higher *gp83* expression may cause the lower Toll-mediated AMP expression, possibly due to lowering *Myd88* expression, which in turn prevents the host from enacting a proper immune response to DiNV infection (Figure 2, Supplementary Figure 5).

**Figure 2:**
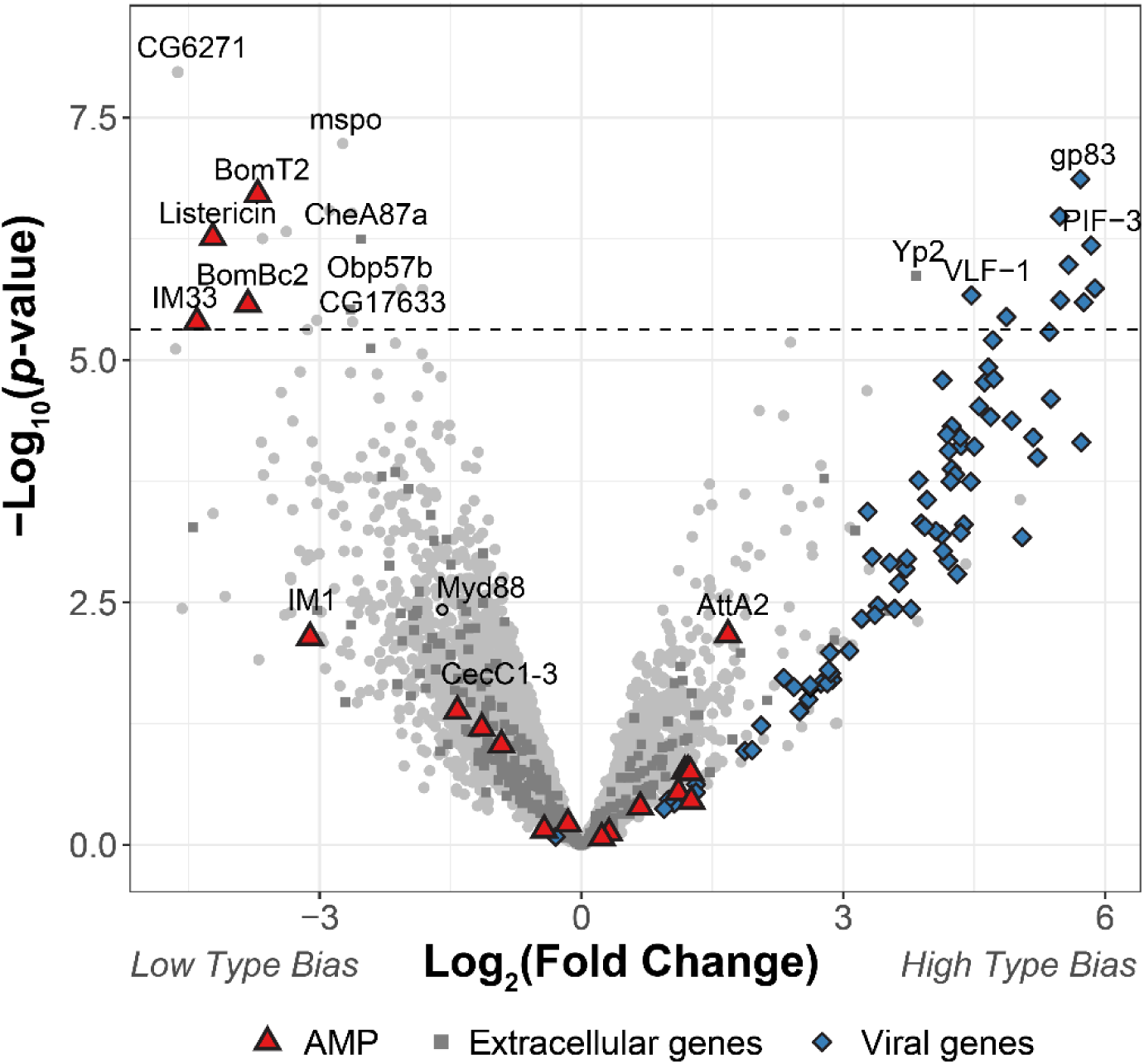
Differential expression of *D. innubila* and DiNV genes between *D. innubila* infected with either the Low type or High type DiNV multilocus genotypes. For host genes, the log-fold change of mRNA fragments per million fragments is compared, while for viral genes the log-fold change of viral mRNA fragments per million fragments per viral particle is compared. Genes are colored/labelled by categories of interest, specifically antimicrobial peptides (AMPs), proteins involved in the extracellular matrix and viral proteins. Specific genes of interest, such as *Myd88*, are also named. The FDR-correct significance cut-off of 0.01 (10,320 tests) is shown as a dashed line.

### Experimental infections recapitulate differences in viral type virulence

To assess if the virulence of the virus types differs in experimental infections, we performed experimental infections of *D. innubila* males using viral filtrate of strains infected with one of the two types of DiNV. As infectious viral titer is increased, survival decreases regardless of viral type (Supplementary Figure 7 & 8, ANOVA residual deviance = 3.536, *p*-value = 2.454e-07, Cox Hazard Ratio z-value > 2.227, *p*-value < 0.02592), with survival decreasing as titer increases (Supplementary Figure 7, Cox Hazard Ratio z-value > 5.428, *p*-value < 5.69e-08). In both cases viral titer also increases for the first 3 days of infection (GLM t-value = 9.817, *p*-value = 3.6e-14).

For a set of independent viral isolates, four High type- and four Low type-infected samples, we diluted samples to roughly equal concentrations of viral particles (about 0.5 particles per host genome prior to filtration) and performed infections for replicates of 10 males with microneedles dipped in one of the filtrate samples. Survival is significantly lower for flies infected with High type viruses when compared to either flies pricked with sterile media (Figure 3A, Cox Hazard Ratio z-value = 3.671 *p*-value = 0.000242) or those pricked with Low type virus (Figure 3A, Cox Hazard Ratio z-value = 4.611 *p*-value = 4e-06). Flies pricked with Low type virus do not show a significant reduction in survival compared to control flies (Cox Hazard Ratio z-value = 1.353, *p*-value = 0.176). We also measured viral titer over time using qPCR, and find titer increases through time in flies infected with either type (Figure 3B, GLM Log_10_(titer) ∼ days + type, days t-value = 9.912, *p*-value = 1.76e-14). Flies infected with High type virus have significantly higher viral titer compared to flies infected with Low type virus (Figure 3B, GLM Log_10_(titer) ∼ days + type, type t-value = 3.934, *p*-value = 0.000211). These results suggest that while the High type strain has higher viral titer and potentially higher transmission rate in wild flies, it also has higher virulence, even after controlling for initial infection titer.

**Figure 3:**
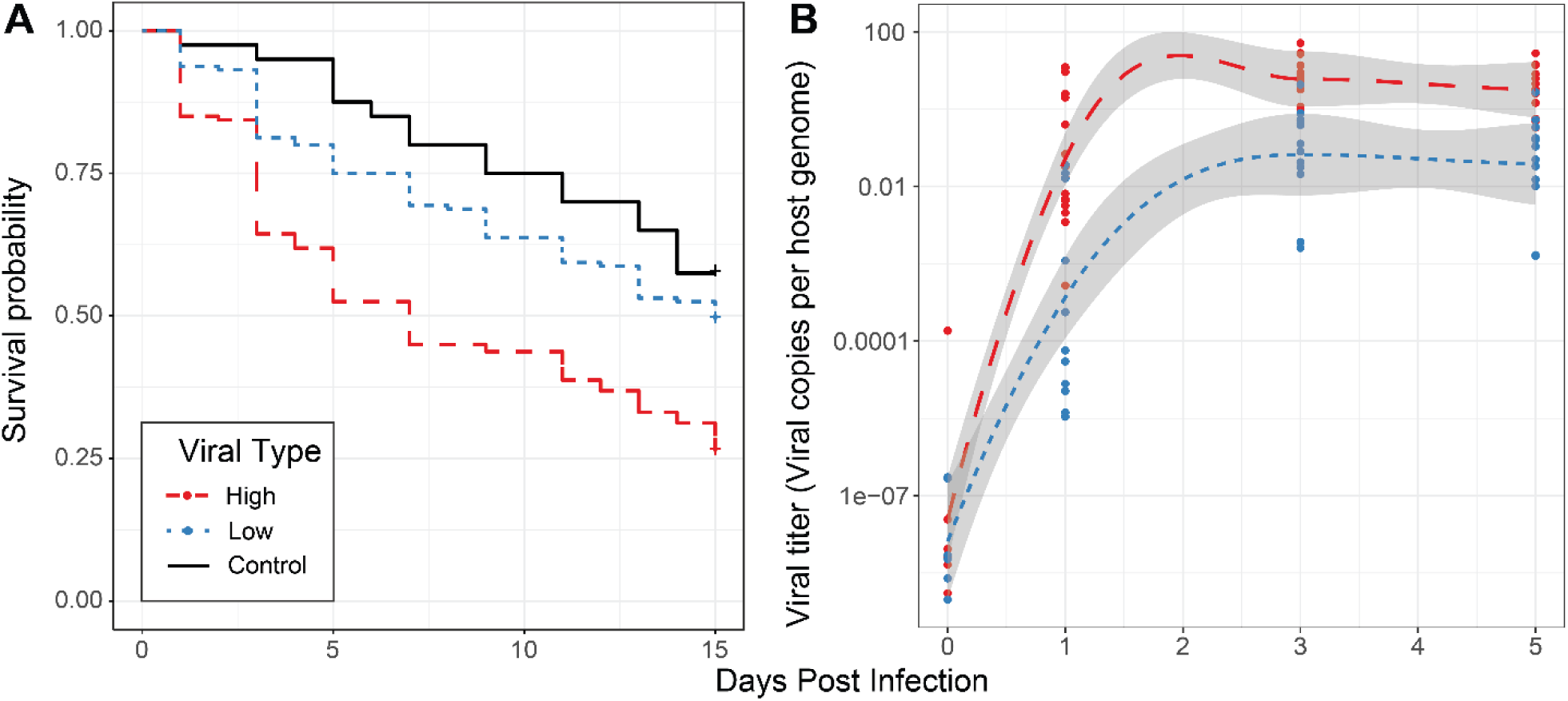
Effect of viral type in experimental infections. **A**. Survival curves of *D. innubila* infected with high and low viral types compared to control flies pricked with sterile media, for 15 days post infection. Survival 5 days post-infection separated by strain is shown in Supplementary Figure 8. **B**. qPCR copy number of viral *p47* relative to *tpi* in *D. innubila* infected with DiNV filtrate of high and low types.

### DiNV types are under strong selection in the host

Recombination is required during nudivirus replication and recombination start sites can be at any point in the single chromosome circular genome (Kelly 1982; Rohrmann 2013). These factors likely cause the incredibly high recombination rates observed in nudiviruses (Blissard and Rohrmann 1990; Wang and Jehle 2009; Rohrmann 2013). In DiNV, the eleven key SNPs that distinguish the High and Low haplotypes are spread across the genome yet are nearly perfectly linked. In contrast, other SNPs in the genome have relatively low linkage disequilibrium, suggesting that selection to maintain each haplotype is strong (Supplementary Figure 4 and 9).

Using McDonald-Kreitman based statistics for detecting selection (Mcdonald and Kreitman 1991; Stoletzki and EYRE-WALKER 2011; Eilertson *et al*. 2012), we tested whether genes that are associated with the High and Low haplotypes exhibited different signatures of natural selection compared to other viral genes. Envelope and virulence proteins show significantly elevated signatures of adaptation (Figure 4, envelope & virulence versus background paired T-test t-value = 2.1761, *p*-value = 0.03814). Genes found in the initial GWAS for virulence, which defined the High and Low types (such as VLF-1, PIF-3, LEF-4 and LEF-5) have significantly higher rate of substitutions being fixed due to selection than background genes (Figure 4, type-associated genes versus all other, t-value = 2.718, *p*-value = 0.00068).

**Figure 4:**
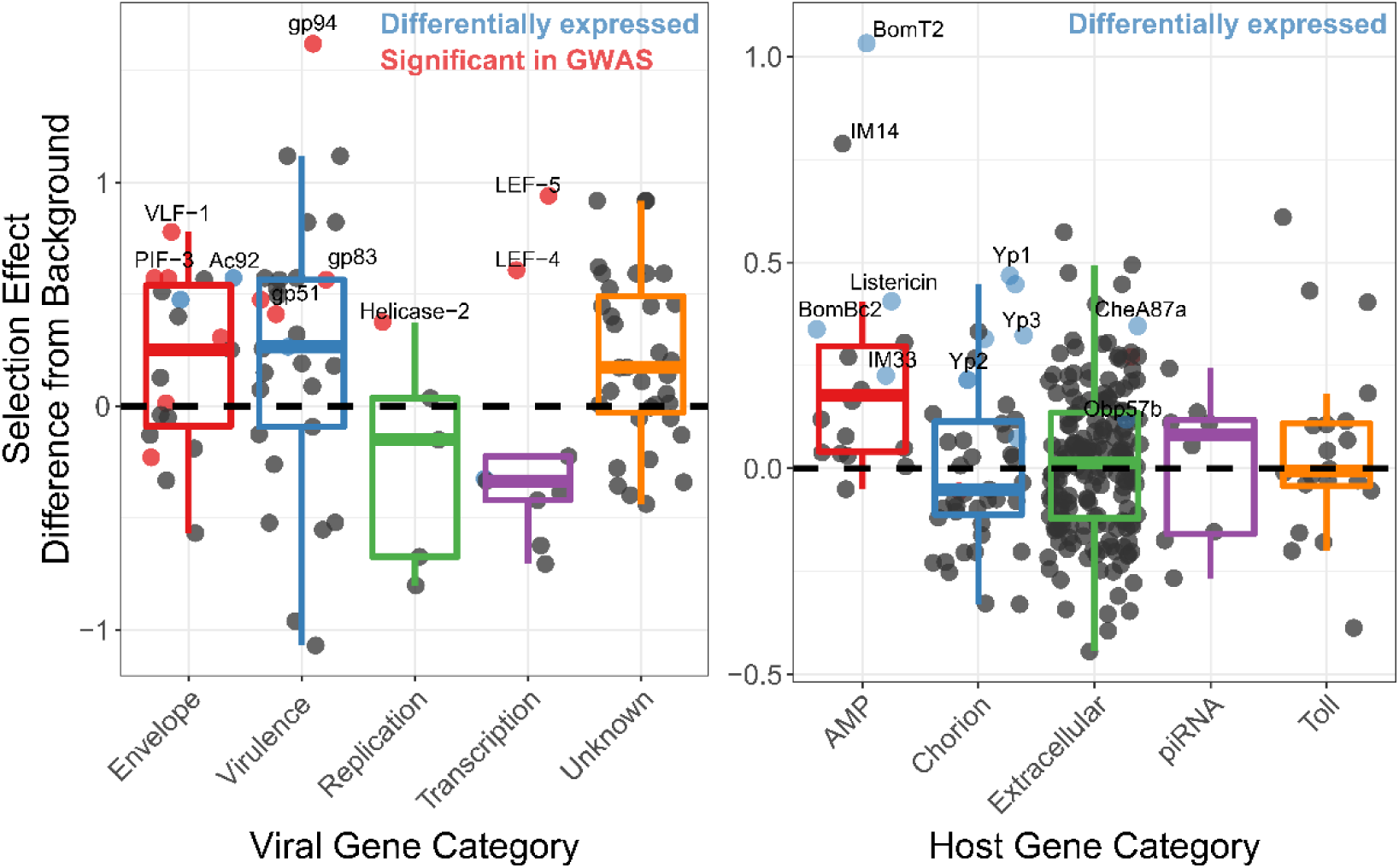
Genes implicated in host/virus interaction are rapidly evolving by positive selection in the Chiricahua population. Difference in selection effect for viral and host gene categories of interest, and nearby background genes, as indicated by the proportion of substitutions fixed by adaptation, weighted by mutations in SnIPRE (Eilertson *et al*. 2012). Genes that have associated SNPs from the GWAS are highlighted in red, while genes which are differentially expressed upon infection, or between viral types are labelled in blue. All GWAS hits are also differentially expressed and labelled in red. Genes of interest are named.

We also performed a GWAS using the host polymorphism and find few associated SNPs, after controlling for the viral haplotype (Supplementary Figure 10). Consistent with the arms race model, host genes we suspect are interacting with DiNV (such as the GWAS hits, AMPs, chorion genes, piRNA genes and extracellular genes) show elevated levels of substitutions fixed by selection compared to background genes in *D. innubila* (Figure 4 & Supplementary Figure 1). Finally, DE chorion genes, extracellular genes and AMPs have significantly more adaptive substitutions than similar non-DE genes (Figure 4, blue dots, differentially expressed versus all other T-test: *D. innubila* t-value = 4.755, *p*-value = 0.000671). Overall these results suggest strong selection is acting on both the host to suppress viral activity and the virus to escape this suppression.

### The High viral type of DiNV evolved repeatedly in three D. innubila populations

We next sought to understand the evolutionary origin of the two types. Given that both types are found in all populations surveyed (Figure 1D), we hypothesized that this could occur one of three ways: First, the derived haplotype was present ancestrally and has been maintained since before geographic isolation occurred. Second, the derived haplotype evolved following geographic isolation and has spread via migration between locations. Third, the derived haplotype has recurrently evolved in each location.

To distinguish between these possibilities and determine the timeframe of divergence, we used the site frequency spectrum of silent DiNV polymorphism to estimate effective population size backwards in time for all populations (Liu and Fu 2015). We find three populations (CH, HU and SR) expand from a single viral particle (N_e_= 1) to millions of particles during the last glacial maximum (30-100 thousand years ago) when *D. innubila* settled its current range (Supplementary Figure 11). This supports a single invasion event during a host-range change. PR appears to expand between 1 and 10 thousand years ago, suggesting a much more recent bottleneck during the range expansion in PR (Supplementary Figure 11) (Hill and Unckless 2020).

We aligned genomic regions containing SNPs to two related nudiviruses, Kallithea virus and Oryctes rhinoceros Nudivirus (OrNV) (Wang *et al*. 2008; Hill and Unckless 2018; Palmer *et al*. 2018b). The High haplotype alleles are not present in either Kallithea or OrNV, and are not found in short read information for wild *D. melanogaster* infected with Kallithea virus (Webster *et al*. 2015), suggesting they are derived in DiNV.

We generated consensus DiNV sequences for each infected *D. innubila* individual and created a whole genome phylogeny to infer geographic diffusion of samples using BEAST (Bouckaert *et al*. 2014). We then performed ancestral reconstruction of the presence of the High type across the phylogeny using APE (Paradis *et al*. 2004). Our samples group as three populations (with HU and SR forming one population) and, consistent with our expectation, the Low type is the ancestral state (Figure 5A). Surprisingly, the High type appears to have evolved repeatedly and convergently within each population, forming separate groups within each population (Figure 5A). The High type also clusters within each population in a principal component analysis of all viral SNPs (Supplementary Figure 9), and when repeating these analyses while excluding the eleven focal SNPs.

**Figure 5:**
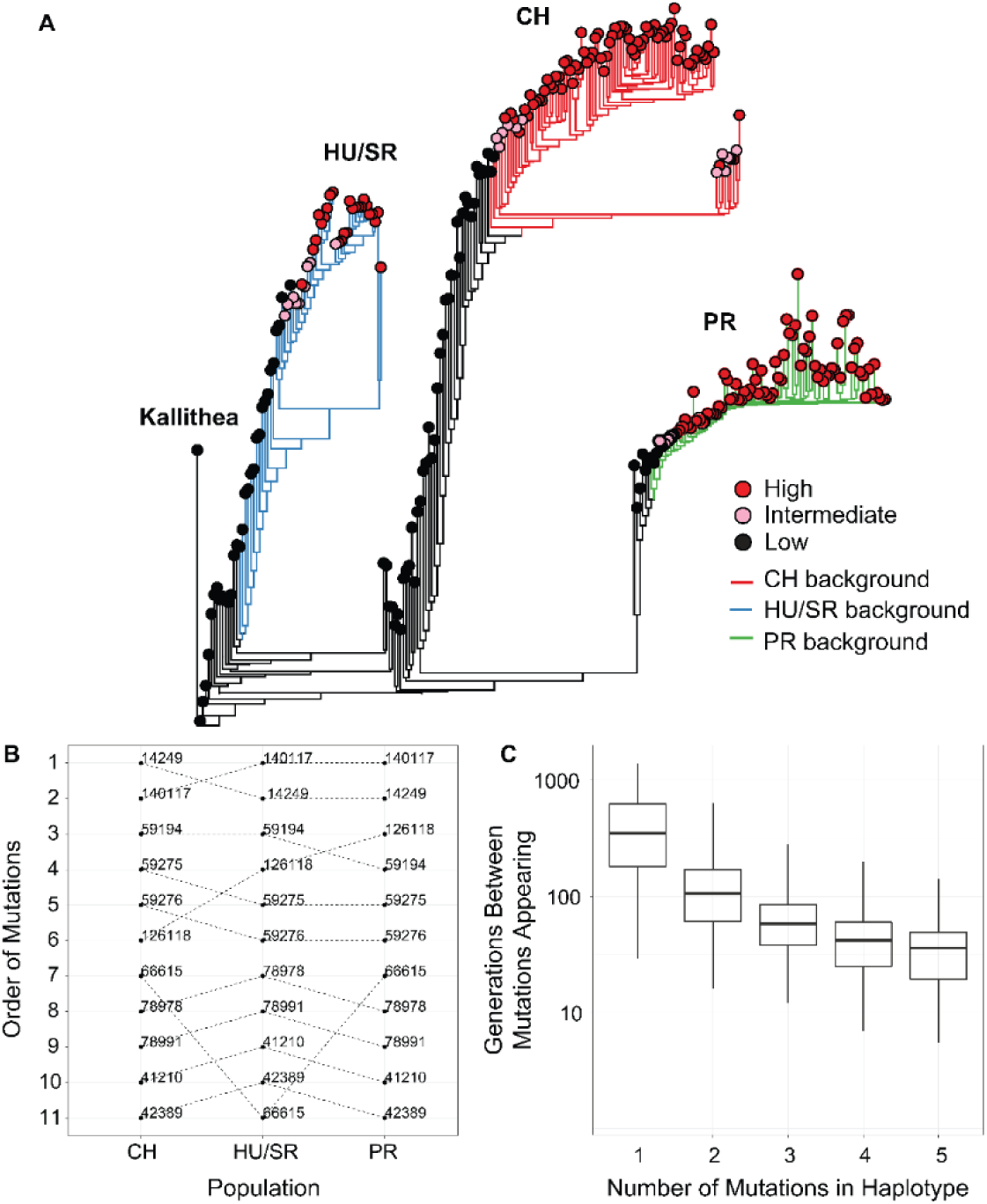
The evolution and maintenance of two viral types. **A**. Phylogeographic reconstruction of the spread of DiNV through *D. innubila*, rooted on the Kallithea virus reference sequence, including a reconstruction of the High type evolution (with strains containing all 11 High type variants shown in red, strains with an intermediate number of high type variants are shown in pink, and strains with no high type variants are shown in black). Branches are colored when the SNPs found in the background for each High haplotype are present in the population, showing that the background differs per population. Black branches show states where branch tips do not contain all the shared High/Low population specific background SNPs. **B**. Order of mutations in the viral haplotype appearing in each population. Apart from three mutations the order is consistent between locations. **C**. The number of generations needed for ‘High titer’ mutations to evolve in simulated populations, given that each mutation increases the mutation rate. The number of generations between each mutation appearing decreases as titer increases.

We next surveyed each background SNP (not associated with the High or Low type) to determine if the general background supports one of the three outlined ways in which the High type evolved and spread in each location. We grouped SNPs by their presence in just the High type or Low type (supporting a single origin and spread by migration) or if they were unique to a single population but shared between the High and Low types (supporting recurrent evolution with recombination).

In total, 391 SNPs (28% of SNPs surveyed) are unique to a single population yet are still shared between both High and Low types (Figure 5 and Supplementary Figure 9), compared to 23 SNPs shared between locations but exclusive to High type samples.

These unique SNPs (161 for CH, 127 for HU and SR, and 53 for PR), are present in all High type samples of a population but a variable proportion of Low types for that population (between 19% and 94%) and are unique to that population. This pattern fits with the High type recurrently evolving on a single background (a different background in each location), supporting recurrent evolution of the High type. The population-specific background SNPs are spread throughout the DiNV genome, with little evidence of recombination with the high type SNPs, making it unlikely that these SNPs recombined onto different backgrounds (Supplementary Figure 4).

Though there is strong linkage between the High type SNPs, they are not perfectly associated with each other (Supplementary Figure 4). Using this slight disassociation and APE (Paradis *et al*. 2004), we performed ancestral reconstruction of SNP origins in each population (assuming recurrent evolution) and find that, excluding three variable SNPs, the evolution of these SNPs was the same order in each population (Figure 5B).

To determine if this recurrent evolution is plausible in our estimated timeframe (∼10,000 years), we simulated viral populations using a modified discrete SIR model using deSolve (Soetaert *et al*. 2010) with estimated baculovirus mutation rates, ranges of viral titer taken our samples and estimated population sizes for each viral population (parameters described in the methods). In this model we used an effective mutation rate scaled to viral titer, considering the mutation rate per particle, so total mutations per generation increase with viral titer. The simulations suggest that waiting time for the first mutation that increases titer is highly variable between replicates but usually occurs within 1000 generations (∼200 years assuming 5 generations per year, in >99.1% of replicates, Figure 5C). The average wait time for each subsequent mutation decreases monotonically (GLM t-value = −2.389, *p*-value = 0.03686). In each case, the next mutation appears in the background of the previous high titer mutation (Figure 5C) due to the elevated effective mutation rate and increased basic reproduction number (R_0_). The accumulation of mutations therefore occurs at a geometric (approximately exponential) rate. Additionally, the standard deviation of time wait times also decreases with each new mutation (GLM t-value = −2.441, *p*-value = 0.04241), increasing the certainty that the entire multilocus genotype will appear in a population rapidly once the initial mutations appear. This chain reaction of adaptation facilitates the repeated evolution of the virulent High type independently in three populations, with all eleven mutations fixing in a population within 6000 generations (∼1200 years) in all replicates (3372 generations on average, ∼675 years), a plausible amount of time given our estimated timeframe.

### Both viral types are found in two other Drosophila species and have also evolved in a geographically distinct population

Since we find two types of DiNV are maintained, and that other species are infected with DiNV, we hypothesized that another species could be a reservoir for the less effective Low type. We chose to study *D. azteca* from the Chiricahuas since it is frequently infected with DiNV (∼33% infection), overlaps with *D. innubila*, and is genetically divergent (40-60 million years) which could mean a very different genetic interaction between host and virus (Unckless 2011). We also examined DiNV-infected *D. falleni* (collected in Georgia) as an outgroup. In all, we sequenced 36 *D. azteca* and 56 *D. falleni*. Both types are present in all examined species, but the high type is rare in *D. azteca* (Figure 6B). The High type has a significantly higher titer than the low type in all cases (Figure 6A). Viral titer is not significantly different across species for either High or Low type (Figure 6A, GLM t-value = −1.351, *p*-value = 0.179). We also find the *D. azteca* samples cluster with CH *D. innubila* samples and contain the CH background SNPs (Supplementary Figure 9C), suggesting no differentiation in the virus infecting different species. Interestingly, *D. falleni* DiNV clusters completely separately from the other samples, likely due to its geographic separation, but still has a derived cluster of High type virus, suggesting a fourth separate evolution of the High type in Georgia. Despite the lack of difference between species samples in Arizona, a lower proportion of the *D. azteca* population is infected with DiNV, and the High Type is less common than the Low type DiNV (Figure 6B). Thus perhaps, even though the relative differences in titer are preserved between the two species, the Low Type is favored in *D. azteca* because this reduced virulence leads to a greater R_0_ in *D. azteca*. Thus, the two types of the virus may be maintained in both host species because though they have become specialized to maximize fitness in one host, messy transmission between host species could lead to their continued presence in both hosts.

**Figure 6:**
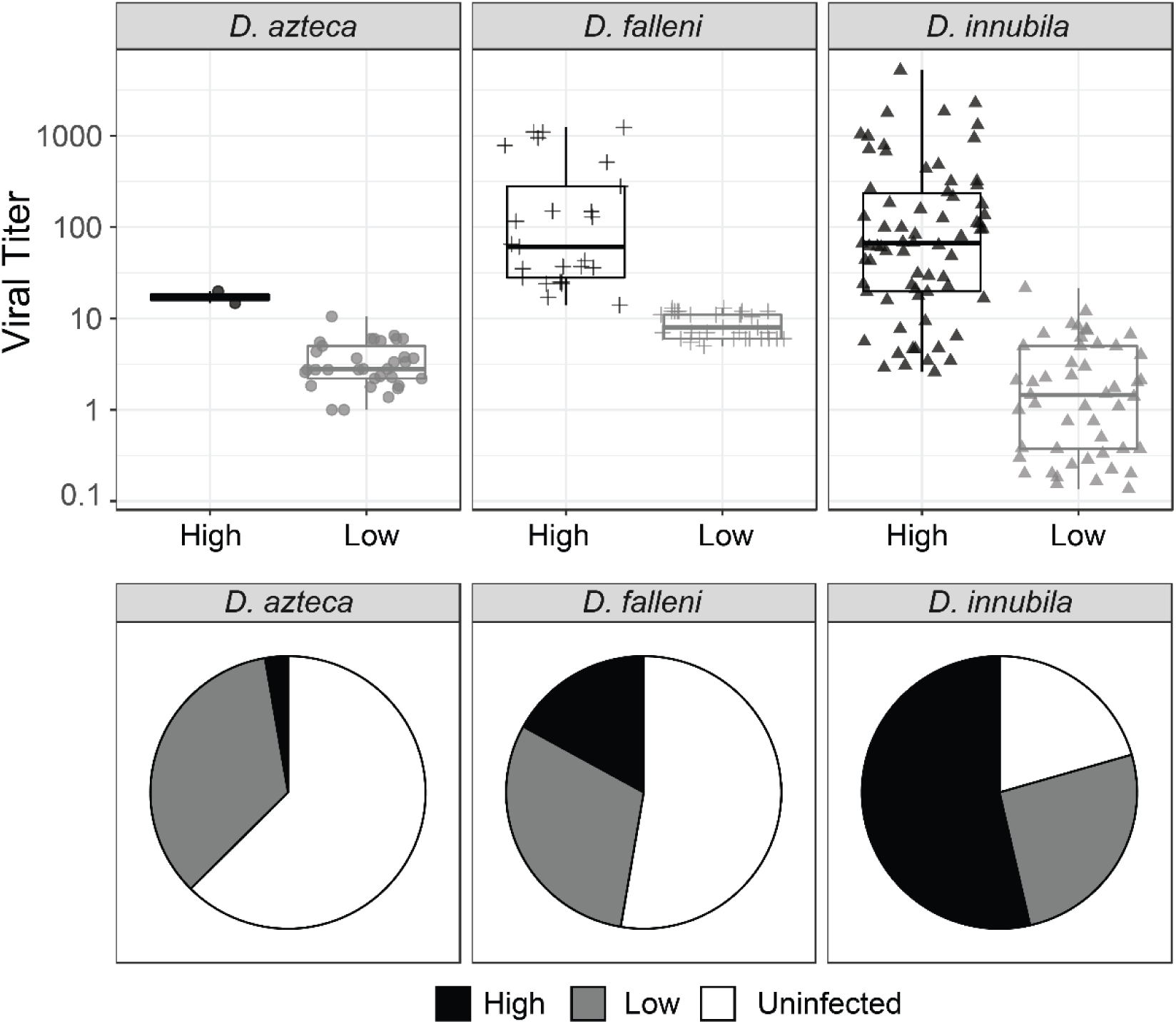
**A**. Viral titer for CH samples of *D. azteca, D. falleni* and *D. innubila* infected with High and Low type DiNV. **B**. Proportion of *D. azteca* and *D. innubila* 2017 CH population infected with High and Low type DiNV.

We also repeated the GWAS for viral titer in DiNV infecting *D. azteca* and *D. falleni*. In both cases we again found the 11 High type SNPs associated with viral titer (GLM t-value > 4.28, *p*-value > 0.0001 in both cases), but not the *Helicase-2* SNP (despite its presence in *D. falleni* DiNV samples). After controlling for the High type, we find no other significant DiNV SNPs in *D. azteca* associated with viral titer. For DiNV infecting *D. falleni*, we find 478 significant SNPs (FDR corrected *p*-value < 0.01), though none of them with as large an effect as the High type associated SNPs.

### Two DiNV types may be maintained due to a trade-off between transmission and virulence

Given the ease that the High type appears to evolve recurrently in populations (Figure 5C), its apparent association with increased infection frequency, and its apparent inability to coinfect with the low type, it is surprising that the High type has not outcompeted the Low type. There are several possible explanations for the maintenance of the two types. First, a soft selective sweep may be occurring on the High type, where recurrent mutation followed by a change in environment changes the fitness of the High type that will eventually result in its fixation (Hermisson and Pennings 2005). Second, both types may be maintained due to a trade-off (Alizon and Van BAALEN 2008). Such a trade-off might even be associated with frequency-dependent selection and cycling frequencies over time. This trade-off could involve different transmission and virulence strategies or might be related to specific adaptation to different hosts.

In a simple model of viral infection dynamics, the success of the virus is measured by its basic reproduction number (R_0_) which is the ratio of the instantaneous transmission rate (β) to the virulence of the virus (γ). If there is a trade-off between transmission and virulence, we might expect that although the High type has a higher instantaneous rate of transmission, it also has higher virulence, killing infected individuals before they can infect other possible hosts. In contrast, those infected with the Low type persist with the infection and can therefore transmit proportionally more virus due to more interactions with susceptible individuals, despite a lower instantaneous transmission rate. To test this, we simulated populations with two viral types using a modified SIR model in deSolve (Soetaert *et al*. 2010). We varied transmission and virulence rates and estimated sets of parameters in which types are maintained within populations. We find that a stable infection frequency depends on both the magnitude of instantaneous transmission rate, and the R_0_, with higher transmission rates increasing the infection frequency, to a maximum of 1 - (γ / β) (Figure 7A). The total infected proportion effectively saturates due to the increased death rate (virulence) of infected individuals, suggesting a trade-off between transmission and virulence as titer increases (Figure 7A). This is consistent with our results in experimental infections in DiNV (Supplementary Figures 7 & 8, Cox Hazard Ratio z-value > 2.227, *p*-value < 0.02592), and other theoretical treatments (May and Nowak 1995; Alizon and Van BAALEN 2008). Two types are only maintained when the R_0_ is equal for each type (γ_1_/β_1_ = γ_2_/β_2_). As the transmission rate of the High type increases, it infects a larger proportion of the population and outcompetes the Low type (with High and Low types at equal proportions when the transmission rates equal), the High type proportion saturates due to the equally increasing virulence rate which keeps the R_0_ equal to the Low type (Figure 7A). Given the requirement for an equal R_0_ for maintenance, the actual proportion of individuals infected with each type depends on the starting infection frequencies of each type and the difference in absolute transmission rate (Figure 7B). Based on the infection frequencies of our sampled populations, we suspect that the High type was able to evolve earlier in the recently bottlenecked PR population (consistent with the High type background being shared with 94% of PR Low types), or the absolute transmission rate (and virulence rate) may have increased in PR population, which is likely what has occurred in CH over time (Figure 7B). Together this suggests that differences in population infection frequencies may depend on a combination of demographic factors, host genetic factors and the instantaneous transmission rate in each population (with lower transmission rates in HU and SR compared to PR and CH). This also implies there is a limit to how virulent a strain can become before it becomes detrimental, as even with higher transmission rates per individual, DiNV may kill the host before it can transmit, reducing its basic reproduction number (R_0_).

**Figure 7:**
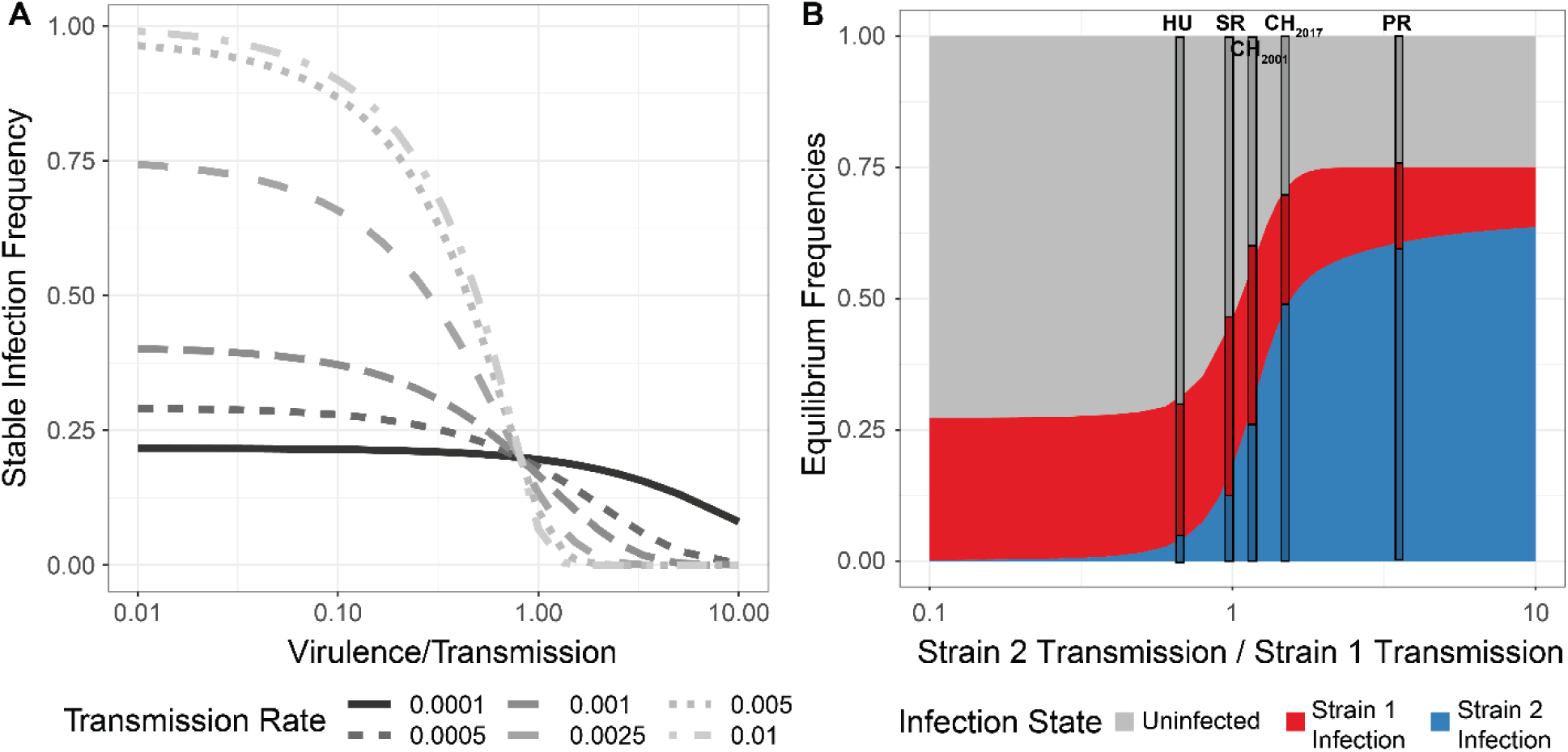
Simulated infected populations to determine parameters of stable infection frequencies of two viral types. **A**. The frequency of stable infection, given the ratio of virulence to transmission, for different magnitudes of transmission but the same starting frequency (0.001%). **B**. Stable infection frequencies given the difference in transmission rates (β) between two viral strains, as strain 2 transmission rate (starting frequency 0.001%) increases relative to strain 1 transmission rate (starting frequency 25%). The difference in infection frequencies of each strain given differing transmission rates of both strains. Stacked bars show the observed frequencies of High type infection (Strain 2, Blue), Low type infection (Strain 1, Red) and Uninfected (Grey) for four sampled populations, positioned on the X-axis to show the estimated ratio of transmission rates for High and Low type, based on the infection frequencies.

## Discussion

Viruses are constantly evolving not just to better infect their host, but also to optimize their infection, to infect as many individuals as possible without preventing the transmission to new hosts (May and Nowak 1995; Lipsitch *et al*. 1996; Alizon and Van BAALEN 2008). Since the host is also evolving in response to the virus, an evolutionary arms-race often ensues (Dawkins and Krebs 1979; Kaltz and Shykoff 1998; Daugherty and Malik 2012). Here, to work towards expanding our understanding of the co-evolution of viruses and their hosts, we examine the population dynamics of Drosophila innubila Nudivirus (DiNV), a DNA virus infecting *D. innubila* (Unckless 2011). DNA viruses have large genomes and often recombination, placing them as a somewhat transitionary pathogen between RNA viruses, bacteria and eukaryotic pathogens and parasites. Within our set of viral samples, we find two DiNV haplotypes which differ by 11 SNPs (Figure 1, named High and Low types). One haplotype (the High type) is associated with higher viral titer, likely due to an increased manipulation of the host immune system and increased expression of viral factors. This derived (High type) has likely recurrently evolved in each population since the last glacial maximum (∼10,000 years ago). The two types appear to be incompatible to some degree, as we find little evidence of co-infections, and mutations appear in a similar order as if navigating an epistatic fitness landscape (Dobzhansky 1937; Kondrashov *et al*. 2002; Gavrilets 2004). Finally, despite the higher titer and transmission rate of the High type, we find that the two types are maintained in all populations sampled, possibly because the increased viral titer also increases the virulence, leading to similar basic reproduction numbers in the High and Low type (Alizon and Van BAALEN 2008).

If the reproduction rate, R_0_, was equal between the two viral types we would expect the infection frequencies to be equal in our sampled populations, which is not the case. This could be caused by the starting frequencies of each type not being equal (with the Low type starting at a higher frequency, Figure 7). There could also be time or host dependent variation, so over time (and between locations), changes in the environment could alter transmission rates of each type, also altering the ratio of High type to Low type. We also do not consider frequency dependent selection in our model, where the transmission rate depends on an interaction between the infection frequencies of the two types.

The R_0_ could be similar between types, but not identical, resulting in slow changes in the ratio of types over time (as seen between 2001 and 2017, Figures 1D & 7). As we only have two time points, we could be witnessing a selective sweep of the High type spreading to fixation (Nielsen 2005), with recombination causing the observed differences in the background. As we find the High type appears to have evolved recurrently, it would be unlikely that we have caught these sweeps partway through in all four populations sampled (Figure 1D). Using the frequency of the High type between 2001 and 2017 CH samples, we can calculate the selection coefficient for the High type if increasing at an exponential rate (which is likely given the intermediate frequency at both time points, Figure 7B). If we assume 5 generations per year and an increase among infected individuals from 39% on 2001 to 71% in 2017, the selection coefficient = 0.007, which suggests the High type would take ∼275 years to fix in a population once it has arisen. Given the coalescence time of the two types is close to the expansion time of the two viruses (2-30 thousand years), this does not fit with our results, suggesting the two types are being maintained and not sweeping. Further, if the High type was sweeping, it would be remarkable for us to catch these sweeps occurring in all four populations sampled, given our estimated time to fixation.

Previous surveys of nudivirus evolution found that a few replication-related genes (including *VLF-1* and *ODV-E56*), likely key targets for host suppression, are under recurrent positive selection (Hill and Unckless 2017; Hill and Unckless 2018). Given that these repeatedly rapidly evolving genes are also associated with the High and Low types and show the highest levels of adaptation (Figure 1, Figure 4), it is entirely possible that these genes are key factors for infection in nudiviruses. They may also be associated with virulence in other nudiviruses or baculoviruses. It is interesting that we find the same SNPs recurrently evolving in each population, opposed to different SNPs affecting the same genes in each population.

One important factor for DNA virus replication is the *Helicase* gene (Blissard and Rohrmann 1990; Rohrmann 2013), and we find variation in *Helicase-2* is associated with viral titer in our survey (Figure 3A & C). As in other nudiviruses, there is extensive adaptive evolution in the *Helicase* gene (Figure 4A) (Hill and Unckless 2017; Hill and Unckless 2018) which has previously been posited to be associated with change in host range (Maeda *et al*. 1993; Croizier *et al*. 1994; Argaud *et al*. 1998; Afonso *et al*. 2001). It is possible the *Helicase-2* variants allow for optimized infection of different species (Croizier *et al*. 1994). In fact, DiNV infects several *Drosophila* species in the New World at varying frequencies (including high prevalence in *D. munda*) (Unckless 2011), and so could have an alternate variant reservoir in any species, with some migration into *D. innubila* (but not *D. azteca* or *D. falleni*), resulting in the appearance of two competing strains. However, this hypothesis ignores the existence of the high and low types which appear to explain the difference in viral titer across multiple species much more convincingly than the *Helicase-2* polymorphism (Supplementary Figure 2).

Nudiviruses have extremely high rates of recombination, as they require at least one crossover during their replication (Rohrmann 2013). Given this high rate of recombination, it is interesting we don’t find more intermediate strains with a mix of High and Low SNPs, supporting the idea of an incompatibility or negative epistatic interactions between SNPs of the two types, similar to a Dobzhansky-Muller incompatibility (Gavrilets 2003).

Some models suggest that viruses are under constant selection to maintain an optimum ratio of virulence to transmission (May and Nowak 1995; Lipsitch *et al*. 1996; Alizon and Van BAALEN 2008). This delicate balance of transmission to virulence could be disrupted with the evolution of a second, high-titer, viral type. In the case of DiNV, the High type has radically higher virulence and appears to compete with the first type impacting the persistence of the virus in a host population. Additionally, the High type appears to recurrently evolve, frequently affecting the persistence of the virus (Figure 5 & 6, Supplementary Figure 9). This posits a situation where the optimum strategy for the ancestral viral type is a reduced transmission rate and the fixation of mutations that are incompatible with High type mutations. Further, this fits with the observed consistent order of fixation for mutations that form the High type (Figure 5B), similar to navigating the neutral adaptive landscape between two incompatible forms in a Bateson-Dobzhansky-Muller incompatibilities (Dobzhansky 1937; Orr 1995; Orr and Turelli 2001; Gavrilets 2003; Orr 2004). This type of viral type interaction is rarely considered in models used for studying infection and could lead to a better understanding of viral dynamics and host-virus co-evolution (Jackson *et al*. 2005; Jackson 2009).

DNA viruses such as DiNV have complicated replication cycles and large genomes. This makes them a sort of evolutionary intermediate between RNA-viruses (small genomes, high mutation rates) and eukaryotes (large genomes, low mutation rates) and tangential to bacteria and archaea (intermediate genomes, low recombination rates). However, adaptation appears to occur through changes in a few key proteins. Here we find the evolution of two competing viral types that differ in these few key genes. These viral types are maintained in populations likely due to a trade-off between transmission and virulence. Overall our results suggest that the high mutation rates and extremely high levels of selection can result in the repeated and convergent evolution of novel host-virus interactions. Additionally, we find that these host-virus interactions for large DNA viruses can be much more complicated than previous models suggest (Dolan *et al*. 2018; Feder *et al*. 2019).

## Materials and Methods

### Fly collection, DNA isolation and sequencing

In this study we used previously collected and sequenced *D. innubila* (Hill and Unckless 2020). Briefly we collected these flies across the four mountainous locations in Arizona between the 22nd of August and the 11th of September 2017. Specifically, we collected at the Southwest research station in the Chiricahua mountains (∼5,400 feet elevation, 31.871 latitude −109.237 longitude), Prescott National Forest (∼7,900 feet elevation, 34.540 latitude −112.469 longitude), Madera Canyon in the Santa Rita mountains (∼4,900 feet elevation, 31.729 latitude −110.881 longitude) and Miller Peak in the Huachuca mountains (∼5,900 feet elevation, 31.632 latitude −110.340 longitude). Baits consisted of store-bought white button mushrooms (*Agaricus bisporus*) placed in large piles about 30cm in diameter, at least 5 baits per location. A sweep net was used to collected flies over the baits in either the early morning or late afternoon between one and three days after the bait was set. Flies were sorted by sex and species at the University of Arizona and were flash frozen at −80°C before being shipped on dry ice to the University of Kansas in Lawrence, KS. During these collections we also obtained *D. azteca* during collections which we also sorted by species and sex and flash froze. *D. falleni* were collected using a similar method in the Smoky Mountains (∼6,600 feet elevation) in Georgia in 2017 by Kelly Dyer, these flies were then sorted at the University of Georgia in Athens GA and shipped on dry ice to the University of Kansas in Lawrence, KS.

For collected *D. falleni* and *D. azteca*, we attempted to assess the frequency of DiNV infection using PCR, looking for amplification of the viral gene *p47*. Using primers from (Unckless 2011), P47F: 5′–TGAAACCAGAATGACATATATAACGC and P47R: 5′–TCGGTTTCTCAATTAACTTGATAGC. We used the following conditions: 95°C 30 seconds, 55°C 30 seconds, 72°C 60 seconds per cycle for 35 cycles.

We sorted 343 *D. innubila* flies, 60 DiNV positive *D. falleni* and 40 DiNV positive *D. azteca* which we then homogenized and used to extract DNA using the Qiagen Gentra Puregene Tissue kit (USA Qiagen Inc., Germantown, MD, USA). We prepared a genomic DNA library of these 343 DNA samples using a modified version of the Nextera DNA library prep kit (∼ 350bp insert size, Illumina Inc., San Diego, CA, USA) meant to conserve reagents. We sequenced the *D. innubila* libraries on two lanes of an Illumina HiSeq 4000 run (150bp paired-end) (Data to be deposited in the SRA). We sequenced the *D. falleni* and *D. azteca* libraries on a separate run of a lane of an Illumina HiSeq 4000 (150bp paired-end).

For 80 male *Drosophila innubila* collected in 2018 (indicated in Supplementary Table 2), we split the sample homogenate in half, isolated DNA from half as described above and isolating RNA using the Direct-zol RNA Microprep protocol (R2061, ZymoResearch, Irvine, CA, USA). We then prepared a cDNA library for each of these 80 RNA samples using a modified version of the Nextera TruSeq library prep kit meant to conserve reagents and sequenced these samples on a NovaSeq NS6K SP 100SE (100bp single end). We also sequenced DNA for these samples, with DNA isolated and prepared as above, also sequenced on a NovaSeq NS6K SP 100SE (100bp single end) (Data to be deposited in the SRA).

### Sample filtering, mapping and alignment

Following sequencing, we removed primer and adapter sequences using cutadapt (Martin 2011) and Scythe (Buffalo 2018) and trimmed all data using Sickle (-t sanger −q 20 −l 50) (Joshi and Fass 2011). We masked the *D. innubila* reference genome (Hill *et al*. 2019), using *D. innubila* TE sequences and RepeatMasker (Smit and Hubley 2008; Smit and Hubley 2013-2015). We then mapped short reads to the masked genome and the Drosophila innubila Nudivirus genome (DiNV) (Hill and Unckless 2018) using BWA MEM (Li and Durbin 2009) and sorted using SAMtools (Li *et al*. 2009). Following this we added read groups, marked and removed sequencing and optical duplicates, and realigned around indels in each mapped BAM file using GATK and Picard (HTTP://BROADINSTITUTE.GITHUB.IO/PICARD; Mckenna *et al*. 2010; Depristo *et al*. 2011). We considered lines to be infected with DiNV if at least 95% of the viral genome is covered to at least 10-fold coverage. We then filtered for low coverage and mis-identified species by removing individuals with low coverage of the *D. innubila* genome (less than 5-fold coverage for 80% of the non-repetitive genome), and individuals we suspected of being misidentified as *D. innubila* following collection. This left us with 318 *D. innubila* wild flies with at least 5-fold coverage across at least 80% of the euchromatic genome, of which 254 are infected with DiNV (Supplementary Table 1). We also checked for read pairs which were split mapped between the DiNV genome and the *D. innubila* genome using SAMtools.

For *D. falleni* we used a previously generated *D. innubila* genome with *D. falleni* variants inserted (Hill *et al*. 2019). We masked the genome with Repeatmasker (Smit and Hubley 2013-2015) and mapped short reads to the masked genome, the repeat sequences and the DiNV genome using BWA MEM and SAMtools (Li and Durbin 2009; Li *et al*. 2009). Then, as with *D. innubila* we filtered for low coverage and mis-identified species by removing individuals with low coverage (less than 5-fold coverage for 80% of the non-repetitive genome) leaving us with 56 *D. falleni* samples infected with DiNV.

For *D. azteca*, we downloaded the genome from NCBI (Accession: GCA_005876895.1) which we then called repeats from with RepeatModeler (Smit and Hubley 2008). We masked the genome with Repeatmasker (Smit and Hubley 2013-2015) and mapped short reads to the masked genome, the repeat sequences and the DiNV genome using BWA MEM and SAMtools (Li and Durbin 2009; Li *et al*. 2009). As with *D. innubila* we then filtered for low coverage and mis-identified species by removing individuals with low coverage of the *D. azteca* genome (less than 5-fold coverage for 80% of the non-repetitive genome), which left us with 37 *D. azteca* samples infected with DiNV. We then called DiNV variation using LoFreq (Wilm *et al*. 2012).

### Calling nucleotide polymorphisms across the population samples

For the 318 sequenced samples with reasonable coverage, for host polymorphism, we used the previously generated multiple strain VCF file, generated using a standard GATK HaplotypeCaller/BCFTools pipeline. We used LoFreq (Wilm *et al*. 2012) to call polymorphic viral SNPs within each of the 254 DiNV infected samples, following filtering using BCFtools to remove sites below a quality of 950 and a frequency less than 5%. We then merged each VCF to create a multiple strain VCF file, containing 5,283 SNPs in the DiNV genome. The LoFreq VCF (Wilm *et al*. 2012) output contains estimates of the frequency of each SNP in DiNV in each sample, to confirm these frequencies, in SAMtools (Li *et al*. 2009) we generated mPileups for each sample and for SNPs of interest (related to viral titer), we counted the number of each nucleotide to confirm the estimated frequencies of these nucleotides at each position in each sample. To confirm that there are no coinfections of types, we also subsampled samples and randomly merged low and high type samples and again generated mPileup files, for SNPs of interest we again counted the number of each nucleotide at each position and confirmed these matched our expected counts in the merged files. We then compared these artificial coinfections to actual samples to confirm the presence or absence of coinfections, finding no samples consistent with coinfections. We then used SNPeff to identify the annotation of each SNP and label synonymous and non-synonymous (Cingolani *et al*. 2012). We extracted the synonymous site frequency spectrum to estimate the effective population size backwards in time using StairwayPlot (Liu and Fu 2015).

### Identifying differentially expressed genes between DiNV infected and uninfected Drosophila innubila

For 100 male *Drosophila innubila* collected in 2018 (indicated in Supplementary Table 2), we homogenized each fly separately in 100µL of PBS. We then split the sample homogenate in half, isolated DNA from half as described above and isolating RNA using the Direct-zol RNA Microprep protocol (R2061). Using the isolated DNA, we tested each sample for DiNV using PCR for *P47* as described previously, using 40 DiNV infected samples and 40 uninfected samples. We then prepared a cDNA library for each of these 80 RNA samples using a modified version of the Nextera TruSeq library prep kit meant to conserve reagents and sequenced these samples on a NovaSeq NS6K SP 100SE (100bp single end). We also sequenced DNA for these samples, with DNA isolated and prepared as above, also sequenced on a NovaSeq NS6K SP 100SE (100bp single end). The DNA sequenced here was mapped as described above, with variation called as described above for other DNA samples.

Following trimming and filtering the data as described in the methods, we mapped all mRNA sequencing data to a database of rRNA (Quast *et al*. 2013) to remove rRNA contaminants. Then we mapped the short read data to the masked *D. innubila* genome and DiNV genome using GSNAP (-N 1 -o sam) (Wu and Nacu 2010). We estimated counts of reads uniquely mapped to *D. innubila* or DiNV genes using HTSEQ (Anders *et al*. 2015) for each sample. Using EdgeR (Robinson *et al*. 2009) we calculated the counts per million (CPM) of each gene in each sample and counted the number of samples with CPM > 1 for each gene. We find that over 70.3% of genes have a CPM > 1 in at least 70 samples. For the remaining genes, we find these genes are expressed in all samples of a subset of the strains (e.g. DiNV uninfected, DiNV infected, DiNV high infected, DiNV low infected). This supports the validity of the annotation of *D. innubila*, given most genes are expressed in some manner, and suggests our RNA sequencing samples show expression results consistent with the original annotation of the *D. innubila* genome.

We attempted to improve the annotation of the *D. innubila* genome to find genes expressed only under infection. We extracted reads that mapped to unannotated portions of the genome and combined these for uninfected samples, samples infected with high type DiNV and samples infected with low type DiNV as three separate samples. We then generated a *de novo* assembly for each of these three groups using Trinity and Velvet (Schulz *et al*. 2012; Haas *et al*. 2013). We then remapped these assemblies to the genome to identify other transcripts and found the consensus of these two for each sample. Using the Cufflinks pipeline (Ghosh and Chan 2016), we mapped reads to the *D. innubila* genome and counted the number of reads mapping to each of these putative novel transcript regions, identifying 15,676 regions of at least 100bp, with at least 1 read mapping in at least 1 sample. Of these, 717 putative genic regions have at least 1 CPM in all 80 samples, or in all samples of one group (DiNV uninfected, DiNV infected, DiNV-low infected, DiNV-high infected). We next attempted to identify if any of these genes are differentially expressed between types, specifically between uninfected strains and DiNV infected strains, and between low-type infected and high-type infected strains. Using a matrix of CPM for each putative transcript region in each sample, we calculated the extent of differential expression between each type using EdgeR (Robinson *et al*. 2009), after removing regions that are under expressed, normalizing data and estimating the dispersion of expression. We find that 26 putative genes are differentially expressed between infected and uninfected types, and 69 putative genes are differentially expressed between high and low types. We took these regions and identified any homology to *D. virilis* transcripts using blastn (Altschul *et al*. 1990). We find annotations for 37 putative genes are either expressed in all samples, or differentially expressed between samples. Of the 14 putative genes expressed in all samples, nine have the closest blast hit to an rRNA gene, and five have hits to unknown genes. For 23 differentially expressed putative genes with blast hits, 3 genes are like antimicrobial peptides (*IM1, IM14, IM3*), these genes are significantly downregulated upon infection, like other Toll regulated AMPs, and have significantly lower expression in strains infected with high type DiNV compared to low types. The remaining 20 genes all have similarity to genes associated with cell cycle regulation, actin regulation and tumor suppression genes.

### Identifying genes associated with viral titer in Drosophila innubila

As the logarithm of viral titer was normally distributed (Shapiro-Wilk test W = 0.05413, *p*-value = 0.342), we used PLINK (Purcell *et al*. 2007) to associate nucleotide polymorphism to logarithm of viral titer in infected samples. We fit a linear model in PLINK including population, sex, *Wolbachia* presence, the date of collection and the relationship matrix for relationship of each sample (inferred using PLINK).

We first fit this model for all 5,283 viral polymorphisms, before performing the association study, we also pruned viral SNPs for both the total population and each subpopulation leaving 1,403 SNPs. For the total sample we identified associations between the logarithm of viral titer and the frequency of the viral polymorphism in each individual sample, resulting in the following model:

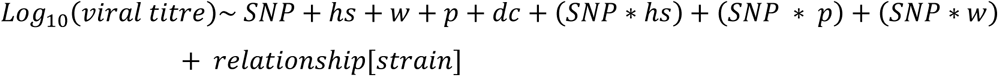

Where hs = host sex, p = location of collection, w = *Wolbachia* presence, dc = date collected Following model fitting, we found factors which seemed to show little or no effect on viral titre (*p*-value > 0.1) using an ANOVA in R (TEAM 2013), and removed these, refitting the model. This was done step-wise, leaving the following model by the end:

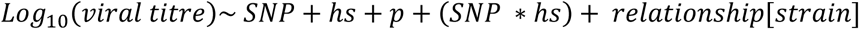

Following this we also performed a GWAS using PLINK (Purcell *et al*. 2007) in the host, using previously called host variation, and considering viral haplotype as an additional covariate.

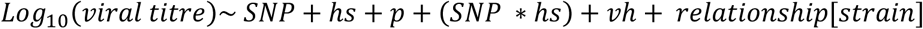

Where hs = host sex, p = location of collection, w = *Wolbachia* presence, dc = date collected, vh = viral haplotype. We found no convincing significant associations (Supplementary Figure 11).

We repeated this analysis for DiNV variants in *D. azteca* and *D. falleni* separately. We performed the GWAS twice, first using the original model, then including viral haplotype as an additional covariate.

### Estimating viral titre using qPCR

Following the identification of the viral haplotype associated with viral titre we sought to determine the effect of viral haplotypes in actual infections. For 20 samples with fly homogenate, we determine the viral titer and haplotype following filtration with a 0.22µM filter.

We performed qPCR for the viral gene *p47* (Forward 5-TCGTGCCGCTAAGCATATAG-3, Reverse 5-AAAGCTACATCTGTGCGAGG-3) on 1 µL of fly filtrate per sample and compared the estimated Cq values across 3 replicates to estimated viral copy number to confirm viral concentration (protocol: 2 minutes at 95°C, 40 cycles of 95°C for 30 seconds and 59°C for 20 seconds, followed by 2 minutes at 72°C). Following this we diluted samples to similar Cq values, relative to the sample with the highest Cq value. We confirmed this by repeating qPCR with *p47* primers of 1 µL of each sample.

For each filtrate sample we performed infections on 30 *D. innubila* males 4-5 days following emergence using pricks with sterile needles dipped in viral filtrate. We recorded survival of each fly each day and removed dead flies. Finally, we took samples 1, 3 and 5-days post infection and measured viral copies of *p47* relative to *tpi* at each time point.

### Phylogeography of DiNV infection and the evolution of the different viral haplotypes

For each DiNV infected *D. innubila* sample, we reconstructed the consensus DiNV genome infecting them using GATK AlternateReferenceMaker and the VCF generated for each strain (Mckenna *et al*. 2010; Depristo *et al*. 2011). We then converted these genomes into Phylip format and used BEAST2 to build the phylogeny of DiNV genomes using 100 million iterations with a burn in of 5 million (Bouckaert *et al*. 2014). We considered phylogeography by providing the longitude and latitude of each samples collection. We then generated a final consensus phylogeny using Tracer with at least 90% majority consensus and removed trees that had not converged to the same joint density (Bouckaert *et al*. 2014). To reconstruct the evolution of the high type viral type we used APE (Paradis *et al*. 2004) to infer the appearance order of the six perfectly linked SNPs on the phylogeny using the all different rates (ARD) discrete model (Paradis *et al*. 2004). We also confirmed these recurrent mutations across the phylogeny using TreeTime to identify recurrently evolving SNPs (Sagulenko *et al*. 2018). Finally, we also created a matrix of SNPs present in at least 2 viral samples and used this matrix in a principle component analysis in R (TEAM 2013), labelling each sample by their viral type in the PCA.

### Simulating the evolution of the high and low viral haplotypes

We sought to simulate the infection of DiNV in *D. innubila* when considering the evolution of a high titer viral haplotype, specifically if two viral types can be maintained against each other at stable frequencies, and if the high viral haplotype with 5 shared mutations could evolve recurrently in the given time period given realistic parameters. We used the R package DeSolve (Soetaert *et al*. 2010) to simulate infection dynamics in a modified SIR model. Specifically, we removed a resistant class, under the assumption that flies won’t live long enough to shed the infection. Therefore, the proportion of population infected per generation is as follows:

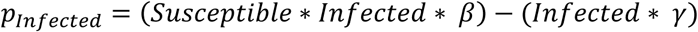

Where *β* represents an infection parameter and *γ* represents a virulence parameter (e.g. the increased likelihood an infected individual has of dying before it can spread its infection). This equation can be rearranged to show the stabilized maximum frequency of infection:

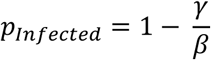

Which is maintained while *γ* is greater than 0.01. The average frequency decreases as absolute transmission rate decreases past this point. We then extended this to include two competing infection types, to represent the two viral types, with the total proportion of population infected per generation as follows:

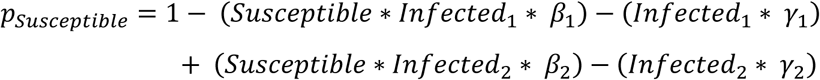

This equation can be rearranged as before to show the stabilized frequency of infection for two viral types:

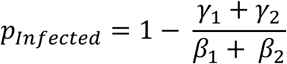

In this case, the two viral haplotypes can both be maintained within a population when the ratio of infection to virulence are the same:

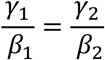

We assessed how starting frequency and difference in *γ* and *β* affects the evolution of each type and their stable frequencies but repeating these simulations for 10,000 generations for each set of parameters. We repeated simulations with transmission rates varying between 0.00001 and 1, and virulence rates set as the transmission rate, divided by a scaling factor between 0.01 and 10 (to vary virulence at multiple rates higher and lower than transmission rate), with all pairwise combinations across all parameters.

Next, we attempted to assess if the ‘high titer’ viral haplotype could possibly evolve recurrently in each population in the time scale seen in our findings. We again used the modified SIR model, this time discrete with population sizes set to 1 million individuals, based on *StairwayPlot* estimates (Liu and Fu 2015). For each infected individual in the population, we tracked the viral titer and also recorded the presence of absence of five mutations, with each mutation increasing the titer of infection, but each further mutation having successively smaller increases in viral titer 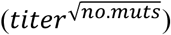, representing the epistatic interaction of high titer associated mutations seen in the viral haplotype. We multiplied the infection parameter, mutation parameter and virulence parameter by viral titer, under the assumption that viral titer increases both infection and death rate, and the mutation rate is per viral particle. We considered a per site mutation rate of 10^−6^, based on estimated baculovirus mutation rate (Rohrmann 2013; Chateigner *et al*. 2015), with a specific mutation rate for the five haplotype mutations of 6.4e-12 (1e-6/155kbp) * viral titer. We then simulated populations in replicate 1000 times for 100,000 generations with a starting infection frequency of 10% for the ‘low titer’ haplotype, recording the frequency of the virus in a population, the frequency of the haplotype and the time that each ‘high titer’ mutation reaches high enough frequency to escape stochastic behavior and behave deterministically under selection (Gillespie 2004).

To estimate the possible selection coefficient for DiNV in the CH population, we assumed an exponential distribution and 5 viral generations per year (80 generations between 2001 and 2017). We then solved the following equation:

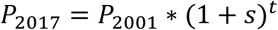

Where P_2017_ = the frequency of the High type among viral samples in 2017 (71%), P_2001_ = the frequency of the High type among viral samples in 2001 (39%), s = the selection coefficient and t = the number of generations (80). We then used this estimated selection coefficient in the same equation to find the number of generations (t) to go from 1/2N_e_s (0.0000714, assuming an N_e_ of 1000000) to fixation (0.99):

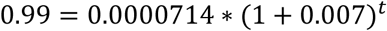

### Experimental infections of Drosophila innubila with DiNV

We chose *D. innubila* samples infected with DiNV and with sequenced genomes, 4 infected with the high type DiNV and 4 infected with the low type. For these samples we estimated their viral copy number per host genome as described previously. We used qPCR on *p47* and *tpi* to find the differences in Cq values to calculate the concentration of each sample relative to the lowest concentration sample and diluted 50µL of filtrate for each sample to match the concentration of each sample to the samples with the lowest titer (IPR07). For a separate 50µL of the IPR01 sample, we performed 1 in 10 serial dilutions to give 45µL of filtrate at full concentration, 1 in 10 concentration, 1 in 100 concentration and 1 in 1000 concentration. Using these sets of samples (matched titer and serial dilutions) we next performed experimental infections.

We transferred 50 *D. innubila* (of roughly equal sex ratio) to new food and let them lay eggs for 1 week, following this we collected male offspring aged 2-5 days for experimental infections.

Across 4 separate days in the mornings (between 9am and 11am), we infected the collected male flies with each sample. For flies in batches of 10, we performed pricks with microneedles dipped in the prepared viral filtrate. For each day we also had 2 control replicates of 10 flies pricked with microneedles dipped in sterile media. Following infections, we checked on each vial of 10 flies one-hour post infection and removed dead flies (likely killed by the needle instead of the virus). We also checked each vial each morning for 15 days, removing dead flies (freezing to determine the viral titer), and flipping flies to new food every 3-4 days. Checking at 10am each day, we recorded the day that each fly died, what filtrate they had been infected with, and what replicate/infection day set they belonged to. We next looked for differences in survival over time compared to sterile wound controls using a Fit proportional hazards regression model in R (TEAM 2013; Kassambara *et al*. 2017), considering titer, infection sample and replicate as co-variates (day of death ∼ [titer or strain] + infection date).

For a second set of experimental infections (performed as described above, stabbed with diluted filtrate from different strains), we also removed 3 living flies 1 hour, 1 day and 5 days post infection. Using qPCR, we found the difference in *p47* log-Cq and *tpi* log-Cq to estimate the viral copy number for each sample over time.

## Acknowledgements

This work was completed with helpful discussion from Justin Blumenstiel, Joanne Chapman, John Kelly, Stuart MacDonald, Andrew Mongue and Carolyn Wessinger. We would especially like to thank Maria Orive, Kelly Dyer and Paul Ginsberg for helpful feedback in the writing of the manuscript and framing of the discussion. Collections were completed with assistance from Todd Schlenke, Paul Ginsberg, Kelly Dyer, Brandon Cooper, John Jaenike and the Southwest Research Station. We thank Brittny Smith and Jenny Hackett at the KU CMADP Genome Sequencing Core (NIH Grant P20 GM103638) and K-INBRE Bioinformatics Core for assistance in genome isolation, library preparation, sequencing and computational resources. This work was supported by a K-INBRE postdoctoral grant to TH (NIH Grant P20 GM103418). This work was also funded by NIH Grants R00 GM114714 and R01 AI139154 to RLU. *D. falleni* collection was funded by NSF grant DEB-1737824.

## Supplementary Figures

**Supplementary Figure 1:**
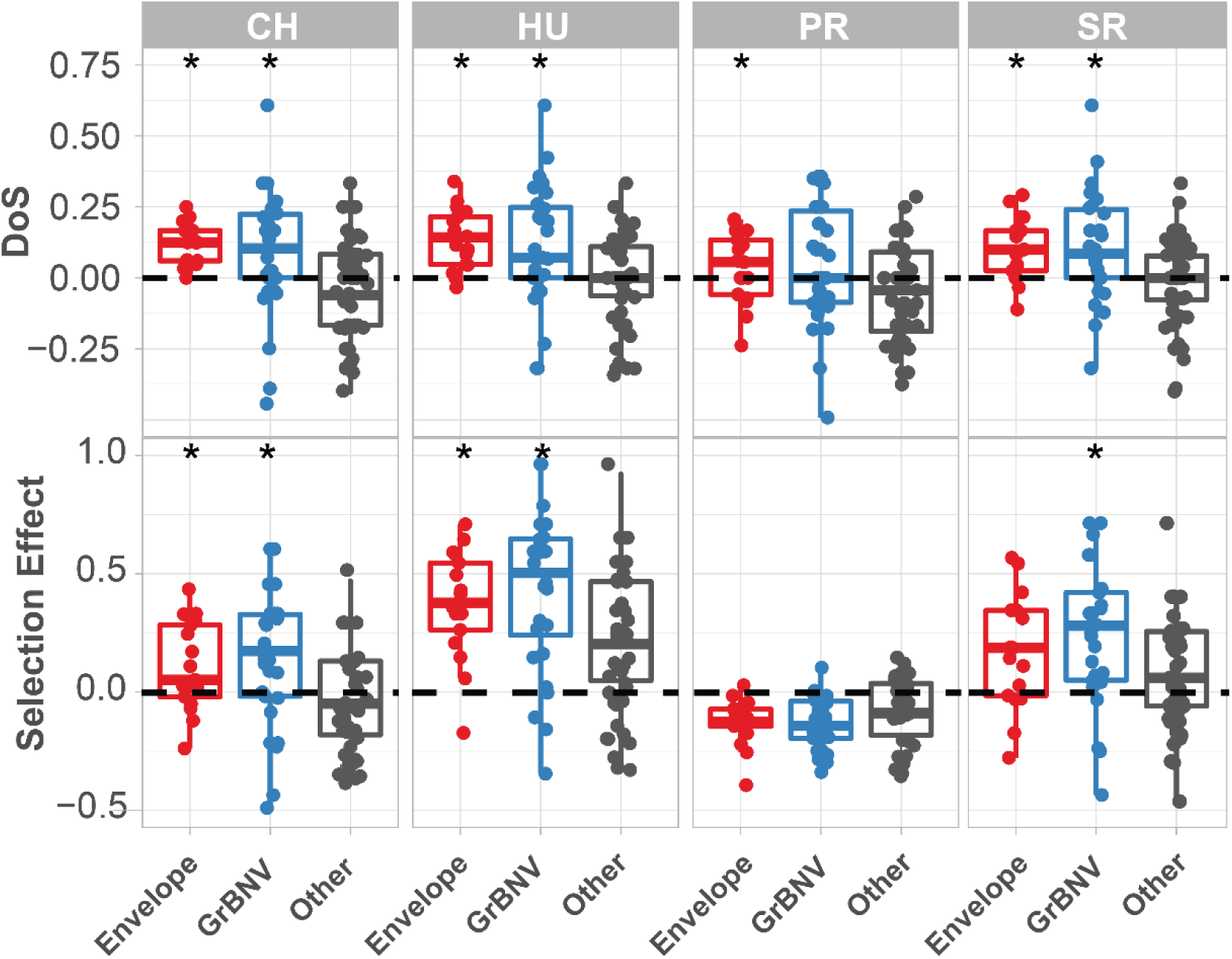
McDonald-Kreitman based statistics for each gene in population of Drosophila innubila Nudivirus, with viral envelope and GrBNV potential virulence factors shown separately. DoS = direction of selection, Selection Effect = SnIPRE estimated weighted DoS. Boxplots marked with a * are significantly higher than background/other viral genes (GLM *p*-value < 0.05).

**Supplementary Figure 2:**
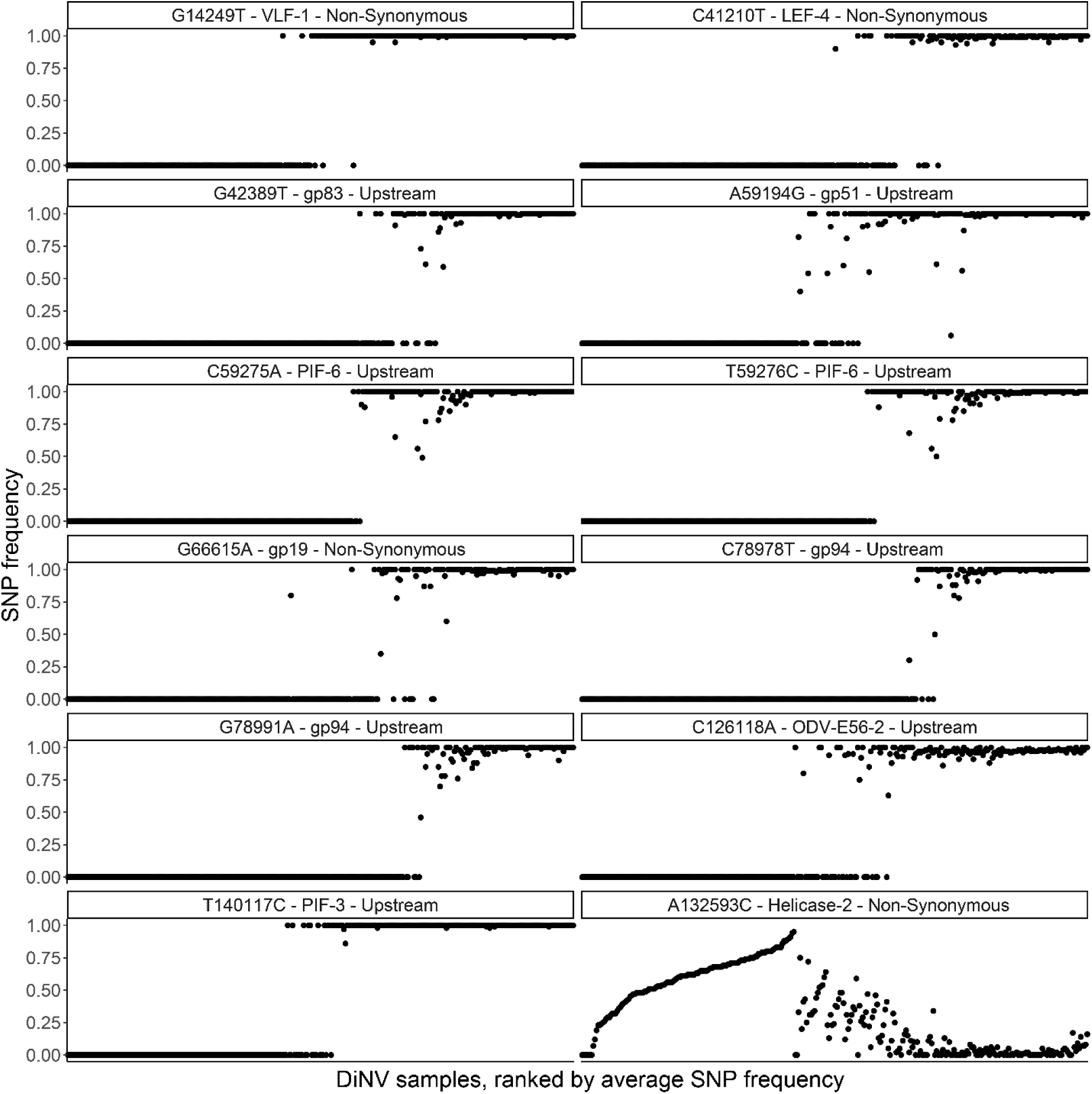
Frequency of each significant SNP within each sample, ranked by the viral titer in each sample (left = lowest, right = highest), to show the strong linkage of SNPs and little evidence of co-infection, also highlights the association between SNP frequency and Helicase-2.

**Supplementary Figure 3:**
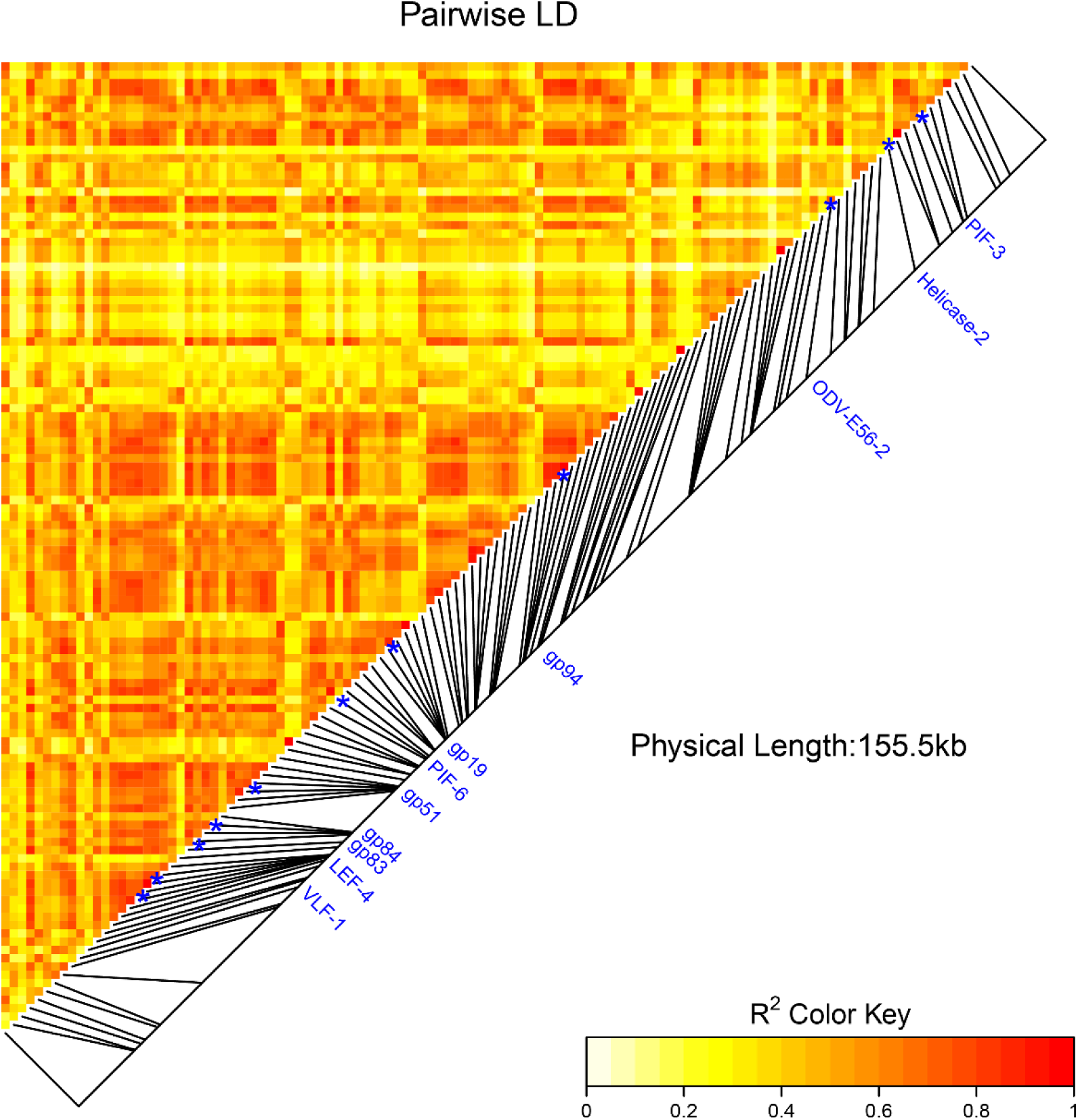
Linkage disequilibrium between SNPs in DiNV. The labelled SNPs (significant SNPs found in the GWAS) are strongly linked. Points are colored by the estimated linkage between SNPs, from red (highly linked, r^2^ = 1) to white (unlinked, r^2^ = 0)

**Supplementary Figure 4:**
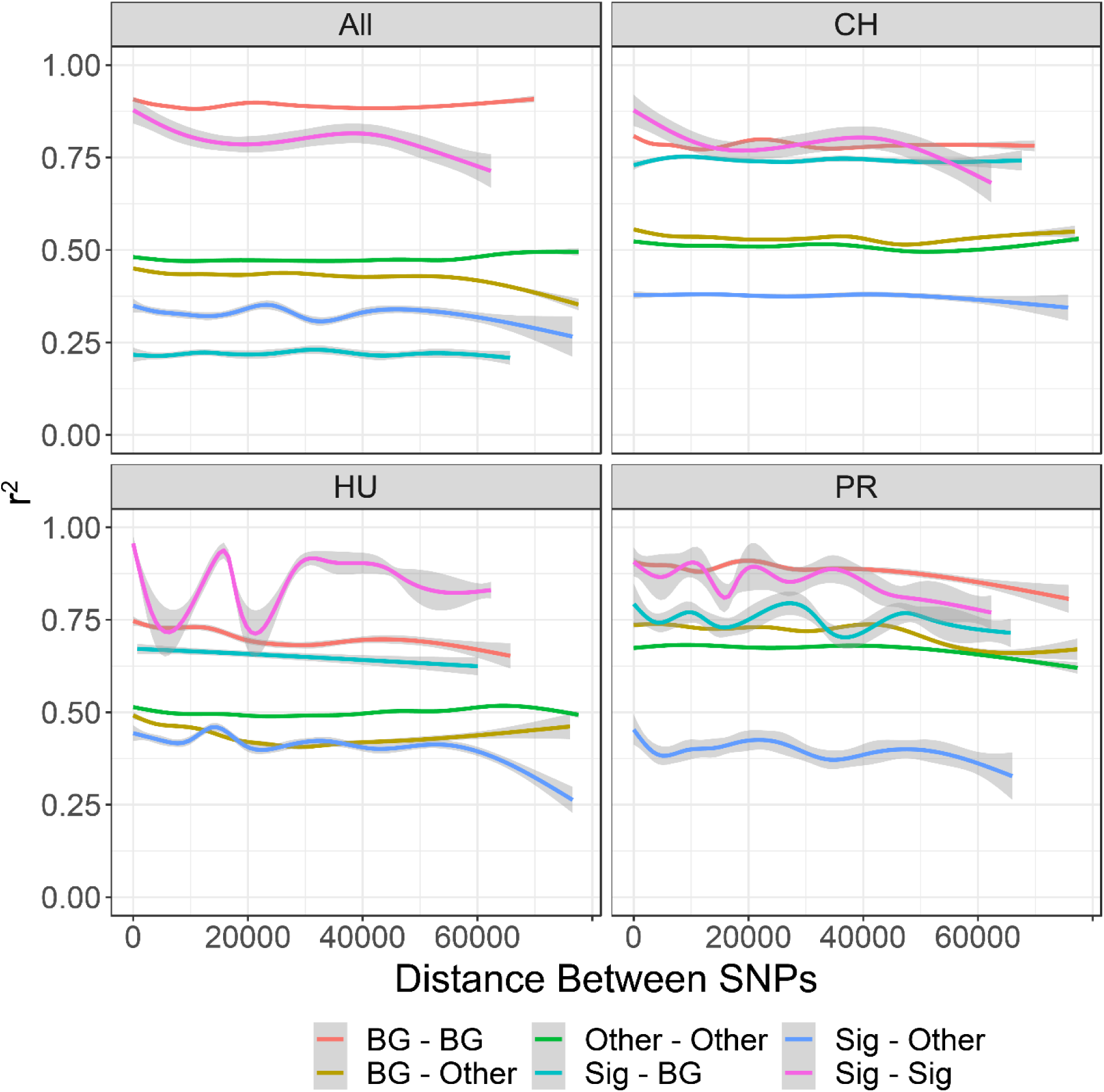
Linkage (r^2^) between different types of SNPs in each population of DiNV, and across all samples. Other = SNPs which are not significantly associated with DiNV titer and do not form the viral haplotype. Sig = SNPs which are significantly associated with DiNV titer and do not form the viral haplotype. BG = SNPs which are in the background which the viral haplotype evolved on in each population.

**Supplementary Figure 5:**
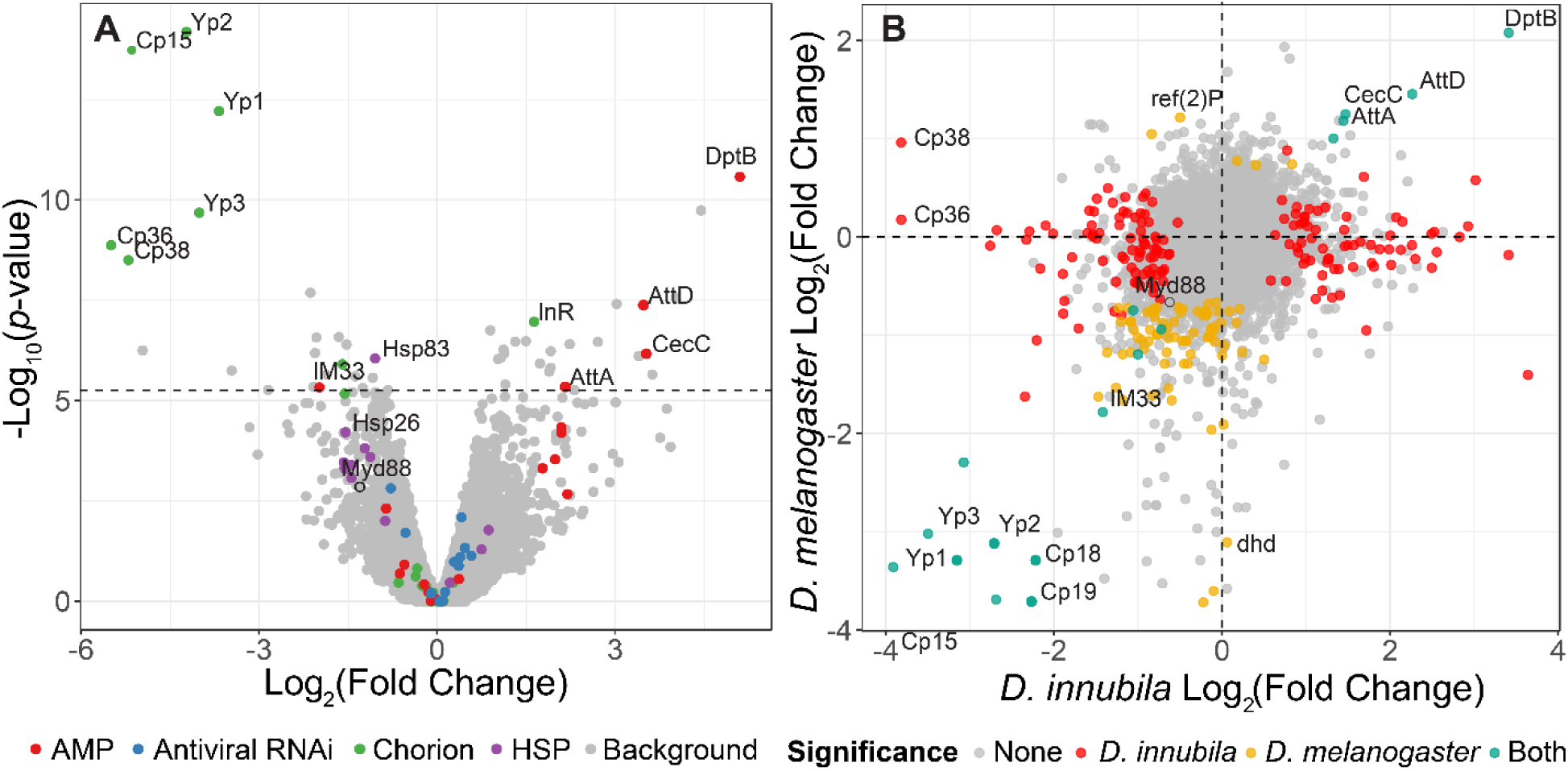
Volcano plot of changes in gene expression between *D. innubila* infected with DiNV and uninfected controls. Gene categories of interest, such as enriched categories, are highlighted in color. The FDR-correct significance cut-off of 0.01 (10,320 tests) is shown as a dashed line. **B**. Comparison of gene expression changes upon infection for *D. innubila* and *D. melanogaster*. Significantly differentially expressed genes (*p*-value < 0.01, FDR-corrected) are colored, genes differentially expressed in both species are colored blue, genes differentially expressed in just *D. melanogaster* are colored yellow and genes differentially expressed in just *D. innubila* are colored red.

**Supplementary Figure 6:**
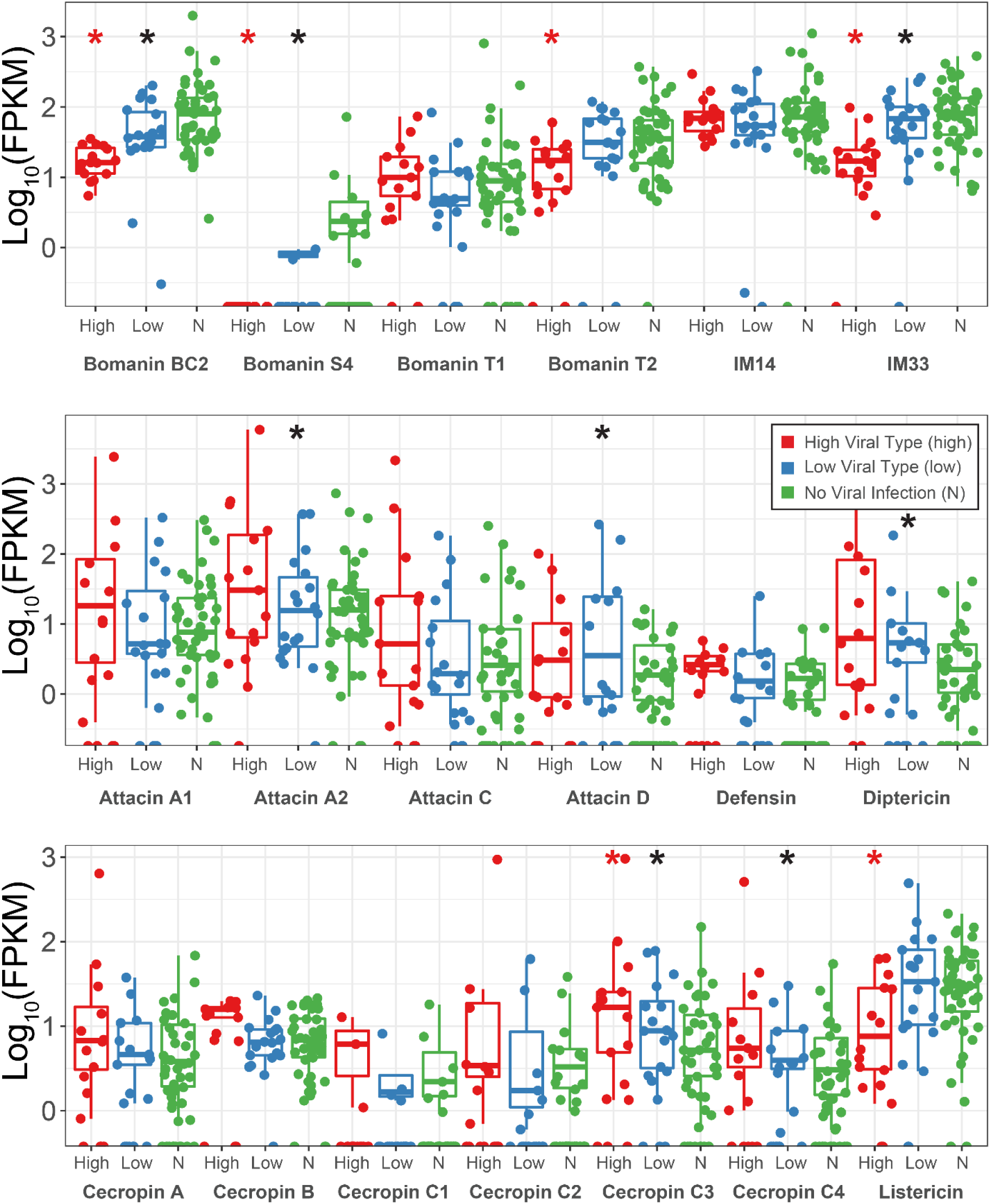
Expression changes (shown as transcript fragments per 1 million reads per 1kbp of exon) of antimicrobial peptides between strains infected with high type DiNV, low type DiNV or not infected. Black stars above low samples show significant differential expression between DiNV infected strains and uninfected strains (multiple testing corrected *p*-value < 0.05). Red stars above high samples show significant differential expression between low type infected strains and high type infected strains (multiple testing corrected *p*-value < 0.05).

**Supplementary Figure 7:**
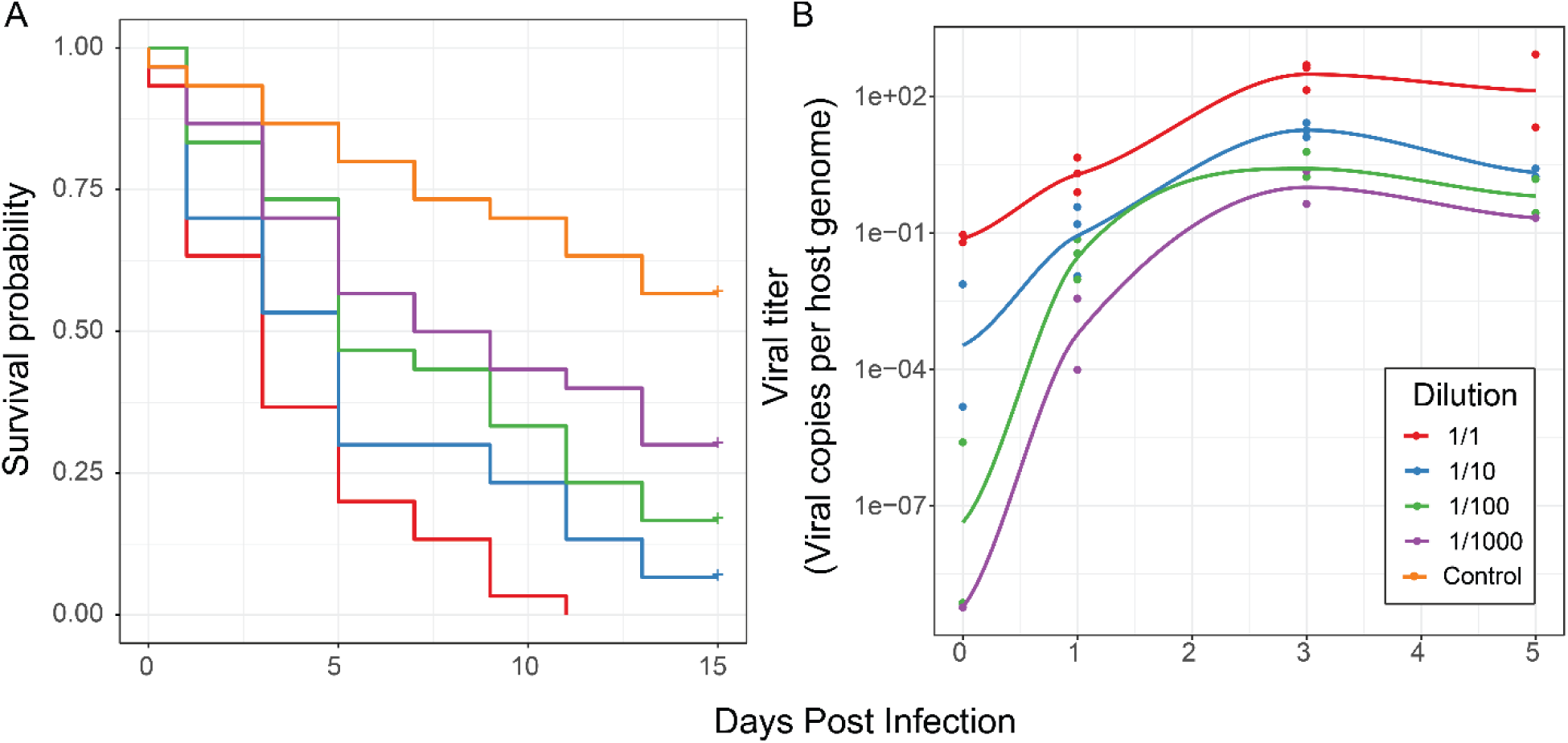
Effect of differences in viral type and titer in experimental infections. **A**. Survival curves of *D. innubila* infected with DiNV filtrate of different dilutions compared to control flies pricked with sterile media, for 15 days post infection. **B**. qPCR copy number of viral *p47* relative to *tpi* in samples of *D. innubila* infected with DiNV filtrate of different dilutions, between 1 and 1000 viral particles per host genome copy.

**Supplementary Figure 8.**
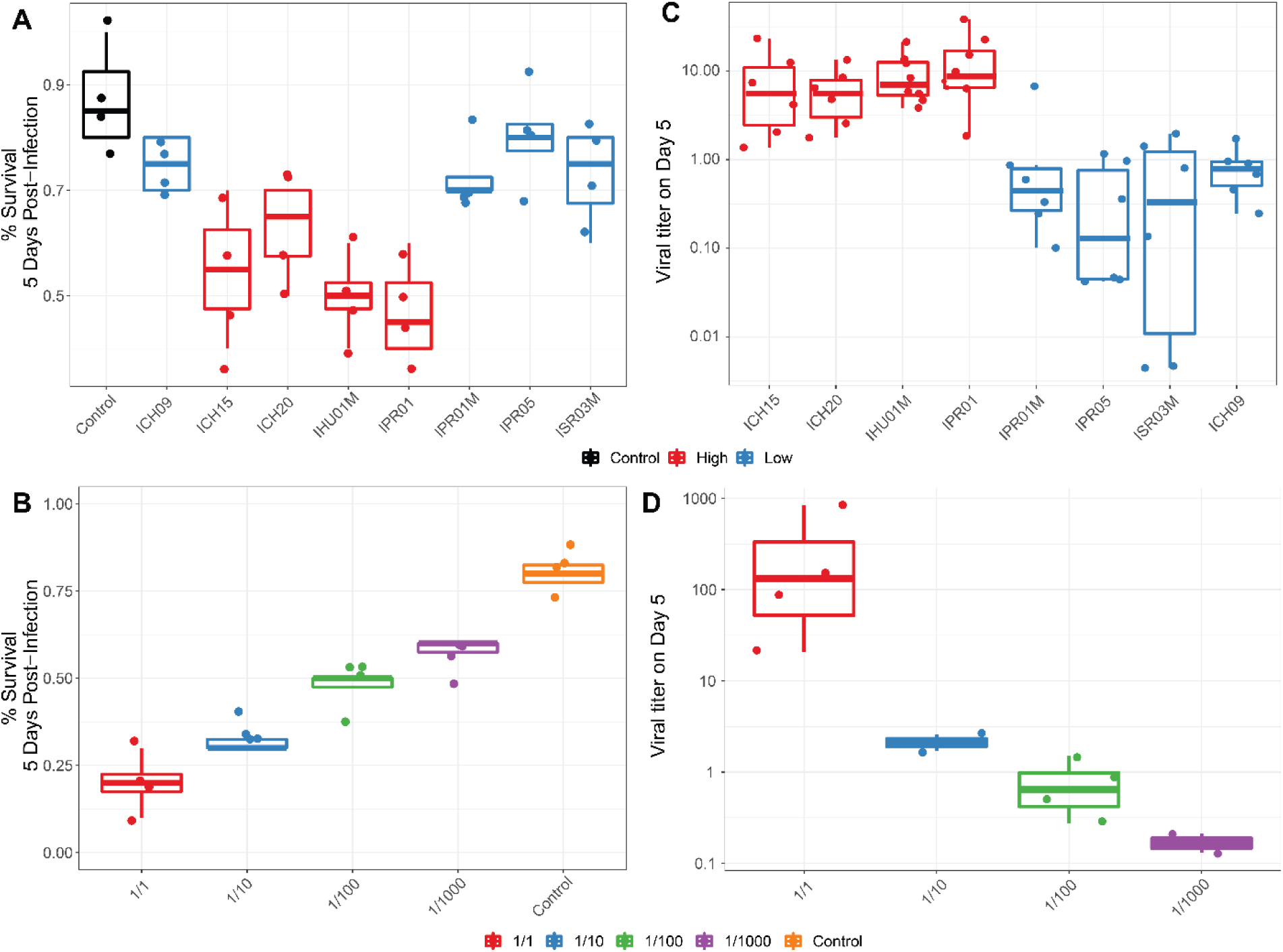
**A:** Survival of *D. innubila* reference strain 5 days post infection, using filtrate from different samples versus uninfected control, colored by high type virus or low type virus. **B**. Survival of *D. innubila* reference strain 5 days post infection using serial dilutions of IPR01 filtrate versus control. **C**. Viral titer estimated per viral genotype at 5 days post-infection, colored by high type virus or low type virus. **D**. Viral titer of DiNV infecting *D. innubila* reference strain 5 days post infection using serial dilutions of IPR01 filtrate versus control.

**Supplementary Figure 9.**
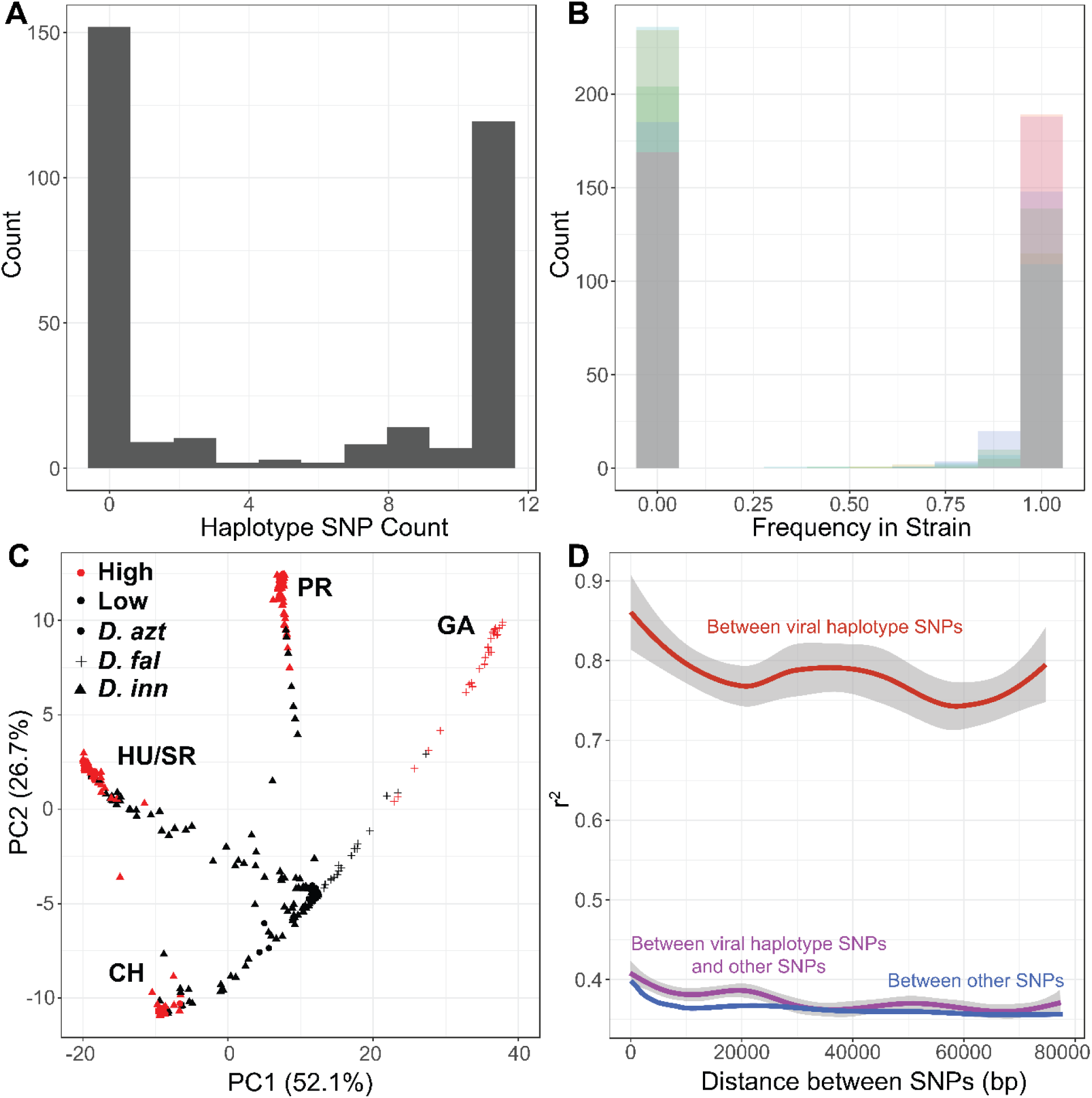
**A**. Frequency of samples with different numbers of SNPs in the viral haplotype, there ae very few intermediate types. **B**. Frequency of each SNP in samples infected with the virus, showing there is little evidence of co-infections. **C**. Principle component analysis of DiNV strains using variation of strains. Strains are colored by the viral type, showing its recurrent evolution. Point shape denotes species in which DiNV was found (*D. azteca, D. falleni or D. innubila*). Strains cluster by collection location. **D**. Linkage between SNPs in the viral haplotype (r^2^) and other SNPs in the haplotype, to other SNPs in the viral genome.

**Supplementary Figure 10.**
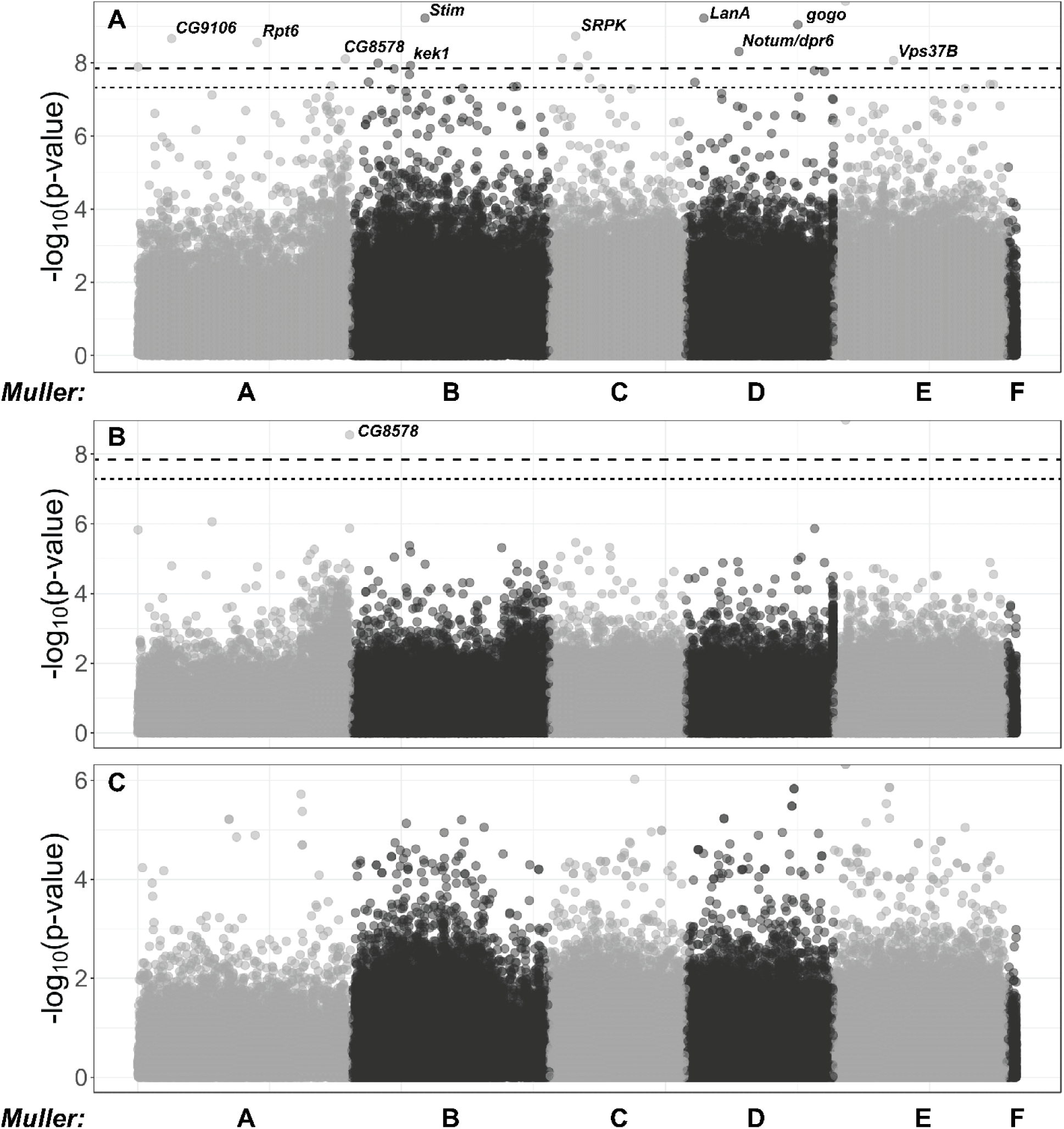
**A:** Manhattan plot of significance of SNP on viral titer after factoring in interaction with the viral haplotype. The significance cut offs are labelled (*p*-value < 0.05 after multiple testing correction dotted, *p*-value < 0.05 after permutations dashed). **B**: Manhattan plot of SNP x viral haplotype interaction for viral titer GWAS in *D. innubila*, calculated using *PLINK*. The significance cut offs are labelled (*p*-value < 0.05 after multiple testing correction dotted, *p*-value < 0.05 after permutations dashed). **C:** Manhattan plot of SNP x sex interaction for viral titer GWAS in *D. innubila*, calculated using *PLINK*.

**Supplementary Figure 11:**
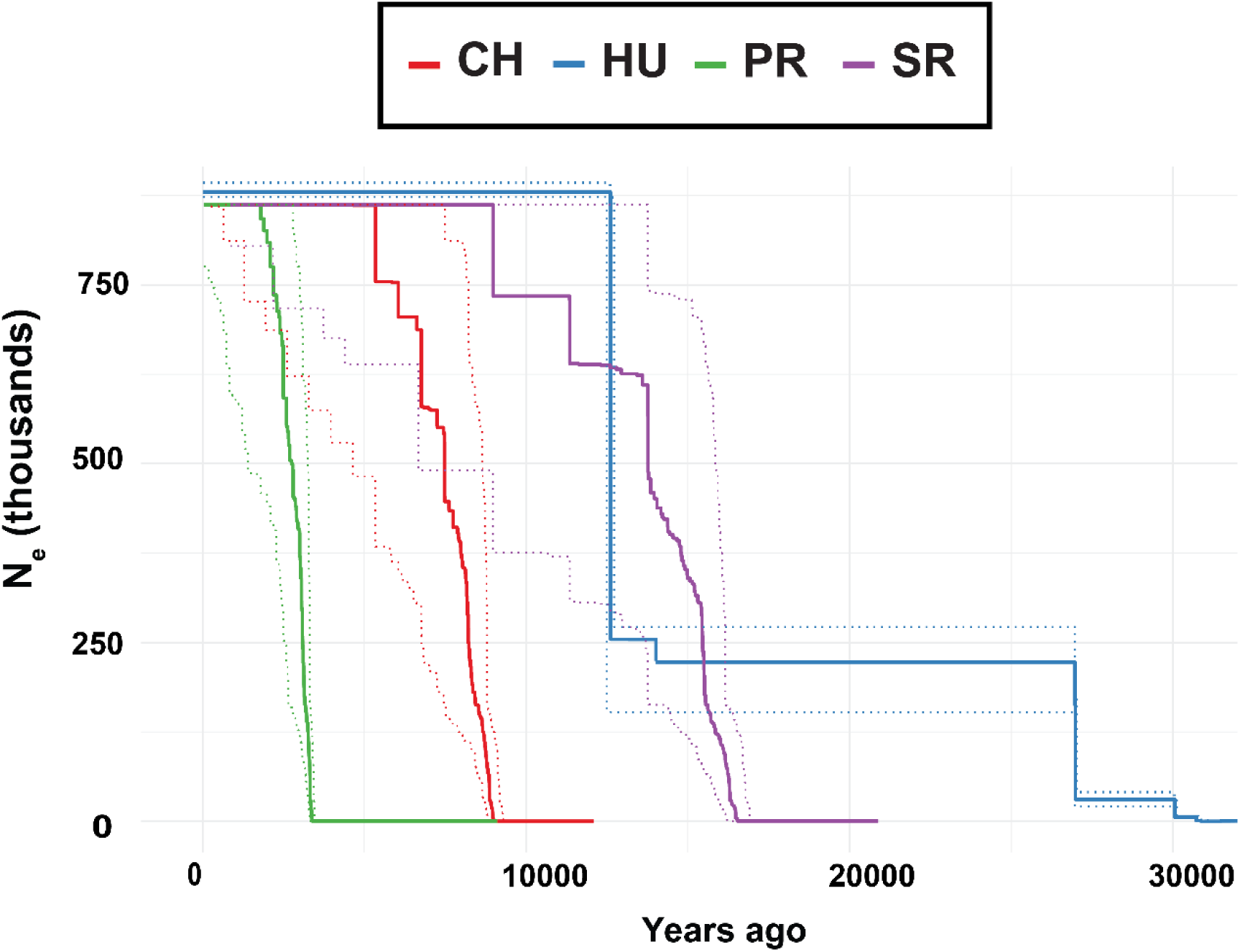
Effective population size backwards for each population of DiNV going backwards in time, estimated using StairwayPlot. Dotted lines indicate the error windows for N_e_ at a given time point. Lines are colored by population.

## Supplementary Tables

**Supplementary Table 1:** Summary of *Drosophila innubila* and *D. azteca* fly samples collected and sequenced for this study, table includes summary of coverage for X chromosome, Muller B, other autosomes, virus and *Wolbachia*. Also contains SRA accessions for each strain.

**Supplementary Table 2:** Summary of *Drosophila innubila* fly RNA and DNA collected and sequenced for this study, including if infected with DiNV.

## Supplementary Data

**Supplementary Data 1:** FPKM of each gene in each sample across the whole *D. innubila* genome, formatted for use in fitting a generalized linear model. Table include the gene name, gene flybase annotation, *D. innubila* name, if the strains is infected with DiNV and the FPKM.

**Supplementary Data 2:** FPKM of each gene in each sample across the whole *D. innubila* genome, formatted for differential gene expression analysis. Table include the gene name, gene flybase annotation, *D. innubila* name, if the strains is infected with DiNV and the FPKM.

**Supplementary Data 3:** Differential gene expression analysis results between viral infected and uninfected strains for both *D. innubila* and *D. melanogaster*. Genes are labelled as if differentially expressed in one of the two species, or if differentially expressed in both.

**Supplementary Data 4:** VCF file for SNPs in DiNV, used in estimation of population genetic statistics and in GWAS using PLINK.

**Supplementary Data 5:** VCF file for SNPs in *D. innubila*, used in estimation of population genetic statistics and in GWAS.

**Supplementary Data 6:** Population genetic statistics calculated for each gene in *D. innubila* using VCFtools for each population.

**Supplementary Data 7:** McDonald-Kreitman statistics calculated for each gene in *D. innubila* using VCFtools for each population.

**Supplementary Data 8:** Population genetic statistics calculated for each gene in DiNV using VCFtools for each population.

**Supplementary Data 9:** McDonald-Kreitman statistics calculated for each gene in DiNV using VCFtools for each population.

